# An Interpreted Atlas of Biosynthetic Gene Clusters from 1000 Fungal Genomes

**DOI:** 10.1101/2020.09.21.307157

**Authors:** Matthew T. Robey, Lindsay K. Caesar, Milton T. Drott, Nancy P. Keller, Neil L. Kelleher

## Abstract

Fungi are prolific producers of natural products, compounds which have had a large societal impact as pharmaceuticals, mycotoxins, and agrochemicals. Despite the availability of over 1000 fungal genomes and several decades of compound discovery efforts from fungi, the biosynthetic gene clusters (BGCs) encoded by these genomes and the associated chemical space have yet to be analyzed systematically. Here we provide detailed annotation and analyses of fungal biosynthetic and chemical space to enable genome mining and discovery of fungal natural products. Using 1037 genomes from species across the fungal kingdom (e.g., Ascomycota, Basidiomycota, and non-Dikarya taxa), 36,399 predicted BGCs were organized into a network of 12,067 gene cluster families (GCFs). Anchoring these GCFs with reference BGCs enabled automated annotation of 2,026 BGCs with predicted metabolite scaffolds. We performed parallel analyses of the chemical repertoire of Fungi, organizing 15,213 fungal compounds into 2,945 molecular families (MFs). The taxonomic landscape of fungal GCFs is largely species-specific, though select families such as the equisetin GCF are present across vast phylogenetic distances with parallel diversifications in the GCF and MF. We compare these fungal datasets with a set of 5,453 bacterial genomes and their BGCs and 9,382 bacterial compounds, revealing dramatic differences between bacterial and fungal biosynthetic logic and chemical space. These genomics and cheminformatics analyses reveal the large extent to which fungal and bacterial sources represent distinct compound reservoirs. With a >10-fold increase in the number of interpreted strains and annotated BGCs, this work better regularizes the biosynthetic potential of fungi for rational compound discovery.

**Significance Statement:** Fungi represent an underexploited resource for new compounds with applications in the pharmaceutical and agriscience industries. Despite the availability of >1000 fungal genomes, our knowledge of the biosynthetic space encoded by these genomes is limited and ad hoc. We present results from systematically organizing the biosynthetic content of 1037 fungal genomes, providing a resource for data-driven genome mining and large-scale comparison of the genetic and molecular repertoires produced in fungi and compare to those present in bacteria.

## Introduction

Fungi have been an invaluable source of bioactive compounds with a wide variety of societal impacts. Mycotoxins such as aflatoxin, ochratoxin, and patulin, pharmaceuticals including penicillin, cyclosporine, and lovastatin, and agrochemicals like paraherquamide and strobilurin are all derived from fungi (1, 2). Recent genome sequencing efforts have revealed that <3% of the biosynthetic space represented by fungal genomes has been linked to metabolite products (3). In both bacteria and fungi, secondary metabolic pathways are typically encoded by biosynthetic gene clusters (BGCs). BGCs encode for backbone enzymes responsible for creating the core metabolite and tailoring enzymes that modify this scaffold, along with regulatory transcription factors and transporters that transport metabolites and necessary precursors (4). In fungi, the most common backbone enzymes include nonribosomal peptide synthetases (NRPSs), polyketide synthases (PKSs), dimethylallyltransferases (DMATs), and terpene synthases.

Over the last decade, genome mining has emerged as an approach that utilizes genome sequencing and bioinformatics for targeted compound discovery based on genes of interest or biosynthetic novelty. Natural products discovery is poised to expand from using a single or few genomes to using many genomes interpreted together using increasingly sophisticated tools (5-9). The interpretation step can infuse knowledge of BGC phylogenetic distribution, inferences about the molecules encoded (*e*.*g*., prevalence and structural variance), and avoidance of known compounds (dereplication). To date, the application of such large-scale genome mining approaches to fungi has been largely limited to individual biosynthetic enzymes (10) or datasets of <100 genomes from well-studied taxonomic groups (11-15).

The concept of a *gene cluster family* (GCF) has emerged as an approach for large-scale analysis of BGCs (5-8). The GCF approach involves comparing BGCs using a series of pairwise distance metrics, then creating families of BGCs by setting an appropriate similarity threshold. This results in a network structure that dramatically reduces the complexity of BGC datasets and enables automated annotation based on experimentally characterized reference BGCs. Depending on the similarity threshold, BGCs within a family are expected to encode identical or similar metabolites and therefore serve as an indicator of new chemical scaffolds. The use of GCFs represents a logical shift from a focus on single genomes of interest to large genomics datasets, providing a means of regularizing collections of BGCs and their encoded chemical space (**Fig. 1A**). The use of GCF networks has been utilized for global analyses of bacterial biosynthetic space (6), bacterial genome mining at the >10,000 genome scale (9, 16), and integrated with metabolomics datasets for large-scale compound and BGC discovery (5, 7). Together with advances to large-scale metabolomics data analysis such as molecular networking (17), the GCF paradigm has helped in the modernization of natural products discovery.

**Figure 1.**
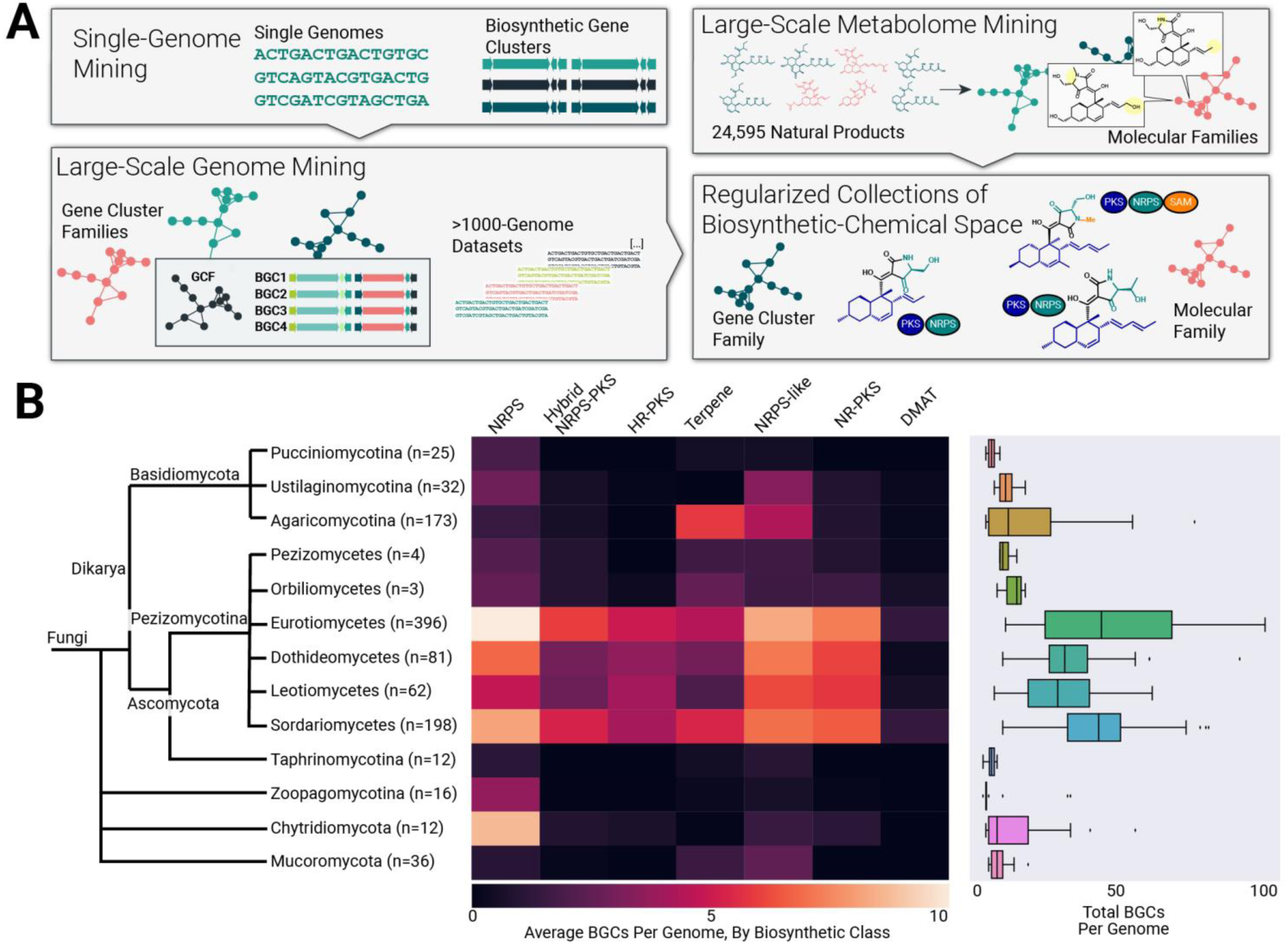
Organizing biosynthetic gene clusters (BGCs) from 1037 fungal genomes. *(A)* Exploring fungal diversity using networks of gene cluster families (GCFs) and molecular families (MFs). A GCF is a collection of similar BGCs aggregated into a network and predicted to use a similar chemical scaffold and create a family of related metabolites. A MF is a collection of metabolites that likewise represent chemical variations around a chemical scaffold. This networking approach enables hierarchical analysis of BGCs and their encoded metabolite scaffolds from large numbers of interpreted genomes. *(B)* Distribution of BGCs across the fungal kingdom. The BGC content of fungal genomes varies dramatically with phylogeny. Organisms within Pezizomycotina have more BGCs per genome and a greater diversity of biosynthetic types than organisms in Basidiomycota and non-Dikarya phyla.

Application of GCFs to fungal genomes has been largely limited to datasets of <100 genomes from well-studied genera such as *Aspergillus, Fusarium*, and *Penicillium* (13-15). Despite the availability of thousands of genomes representing a broad sampling of the fungal kingdom, global analyses of the BGC content of these genomes are lacking. As such, our knowledge of the overall phylogenetic distribution of GCFs in fungi is limited, and many taxonomic groups have no experimentally characterized BGCs. Therefore, we performed a global analysis of BGCs and their families from a dataset of 1037 genomes from across the fungal kingdom. Across Fungi, the vast majority of GCFs are species-specific, indicating that species-level sampling for genome sequencing and metabolomics will yield significant returns for natural products discovery.

To relate this now-available set of fungal GCF-encoded metabolites to known fungal scaffolds, we performed network analysis of 15,213 fungal compounds, organizing these into 2,945 molecular families (MFs) (**Fig. 1A**). Analysis of this joint genomic-chemical space revealed dramatic differences between both major fungal taxonomic groups, as well as between bacteria versus fungi. This lays the groundwork for systematic discovery of new compounds and their BGCs from the fungal kingdom.

## Results

### A reference set of fungal biosynthetic gene clusters

Despite the availability of thousands of fungal genomes, the biosynthetic space represented within them has yet to be surveyed systematically. To address this gap, we curated a dataset of 1037 fungal genomes, covering a broad phylogenetic swath (**Table S1**; **Dataset S1)**. This selection includes well-studied taxonomic groups such as Eurotiomycetes (*Aspergillus* and *Penicillium* genera) and Sordariomycetes (*Fusarium, Cordyceps*, and *Beauveria* genera), and groups for which we have little knowledge about their BGCs, such as Basidiomycota or Mucoromycota. This genomic sampling likewise covers a large swath of ecological niches, from forest-dwelling mushrooms to plant endophytes to extremophiles (18).

Each of the 1037 genomes was analyzed using antiSMASH (19), yielding an output of 36,399 BGCs ranging from 5 to 220 kb in length. As has been previously observed (20), the number of BGCs per genome varies dramatically across Fungi (**Fig. 1B**; **Table S1**). Eurotiomycetes average 48 BGCs per genome, with 25% of organisms within this class possessing >60 BGCs. Organisms outside of Pezizomycotina possess significantly fewer BGCs, with organisms from the non-Dikarya phyla averaging <15 BGCs per genome. The distribution of biosynthetic classes across the fungal kingdom also varies dramatically and unexpectedly. Organisms within the Pezizomycotina classes Eurotiomycetes, Dothideomycetes, Leotiomycetes, and Sordariomycetes average approximately 5 each of NRPS, hybrid NRPS-PKS, NRPS, HR-PKS, terpene, NRPS-like, and NR-PKS, and 2 DMAT BGCs per genome (see **Fig. 1B**). Basidiomycota have far fewer BGCs encoding a relatively limited chemical repertoire, with terpene BGCs being the most abundant in Agaricomycotina as previously implied (10).

### Organizing gene clusters into families to map fungal biosynthetic potential

To further assess the ability of fungi to produce new chemical scaffolds, we grouped BGCs into families using the pairwise distance between BGCs and a clustering algorithm to yield GCFs. BGCs from antiSMASH were converted to arrays of protein domains then compared based on the fraction of shared domains and backbone protein domain sequence identity (7, 8). DBSCAN clustering was performed on the resulting distance matrix, resulting in a set of 12,067 GCFs (**Fig. 2A**) organized into a network (**Fig. 3A**). Across the fungal kingdom, the distribution of GCFs shows a clear relationship with phylogeny (see yellow streaks in **Fig. 2A, Figs. S1-S5**). In isolated studies of well-characterized strain sets of *Aspergillus* and *Penicillium*, GCFs have been thought to be largely genus- or species-specific (13, 21, 22); however, here we show that several GCFs span entire subphyla or classes (**Fig. 2A**). The fraction of GCFs that two organisms share is likewise correlated with phylogenetic distance, evidenced by sets of shared GCFs between closely related taxonomic groups (**Fig. S6**). In order to facilitate visualization of these phylogenetic patterns, we created *Prospect*, a web-based application for hierarchical browsing of GCFs, BGCs, protein domains and annotations for known compound/BGC pairs (http://prospect-fungi.com). Additional details of the site are available in **SI Methods**.

**Figure 2.**
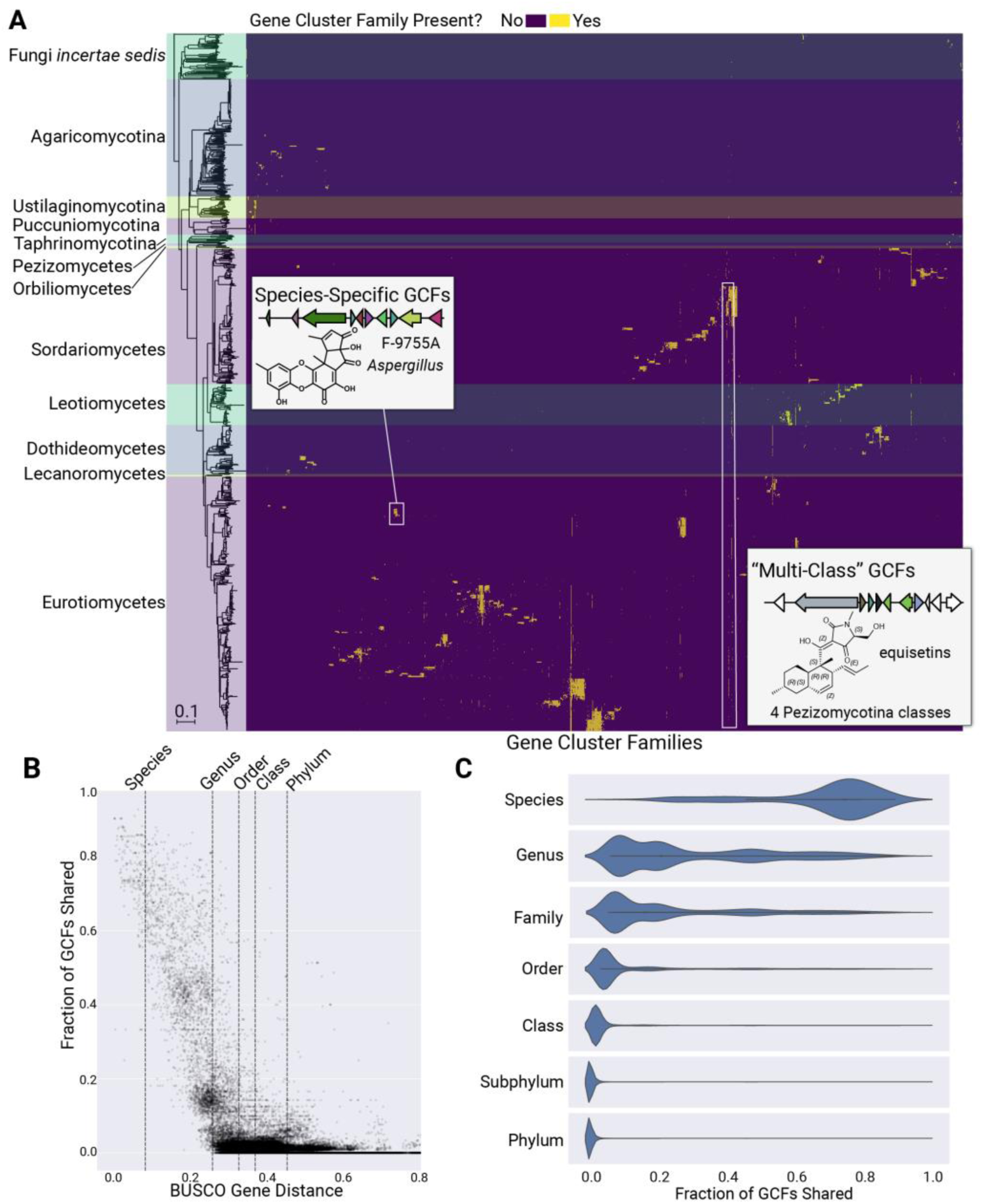
The distribution of 12,067 gene cluster families (GCFs) across the fungal kingdom. *(A)* Heatmap of GCFs across Fungi. The phylogram to the left shows a Neighbor Joining species tree based on 290 shared orthologous genes across 1037 genomes; horizontal shaded regions across the heatmap correspond to each labeled taxonomic group. The order of GCF columns is the result of hierarchical clustering based on the GCF presence/absence matrix. Across Fungi, the distribution of GCFs largely follows phylogenetic trends, with most GCFs confined to a specific genus or species. *(B)* Relationship between genetic distance and GCF content. The dotted lines indicate median genetic distance values for organisms within the same species, genus, order, class, or phylum. Each point in the scatterplot represents a pair of genomes and the fraction of the pair’s GCFs that are shared. *(C)* Relationship between taxonomic rank and shared GCF content across the fungal kingdom. Violin plots show the fraction of GCFs shared between all pairs of organisms within our 1000-genome dataset, with each pair classified based on the lowest taxonomic rank shared between the two organisms.

**Figure 3.**
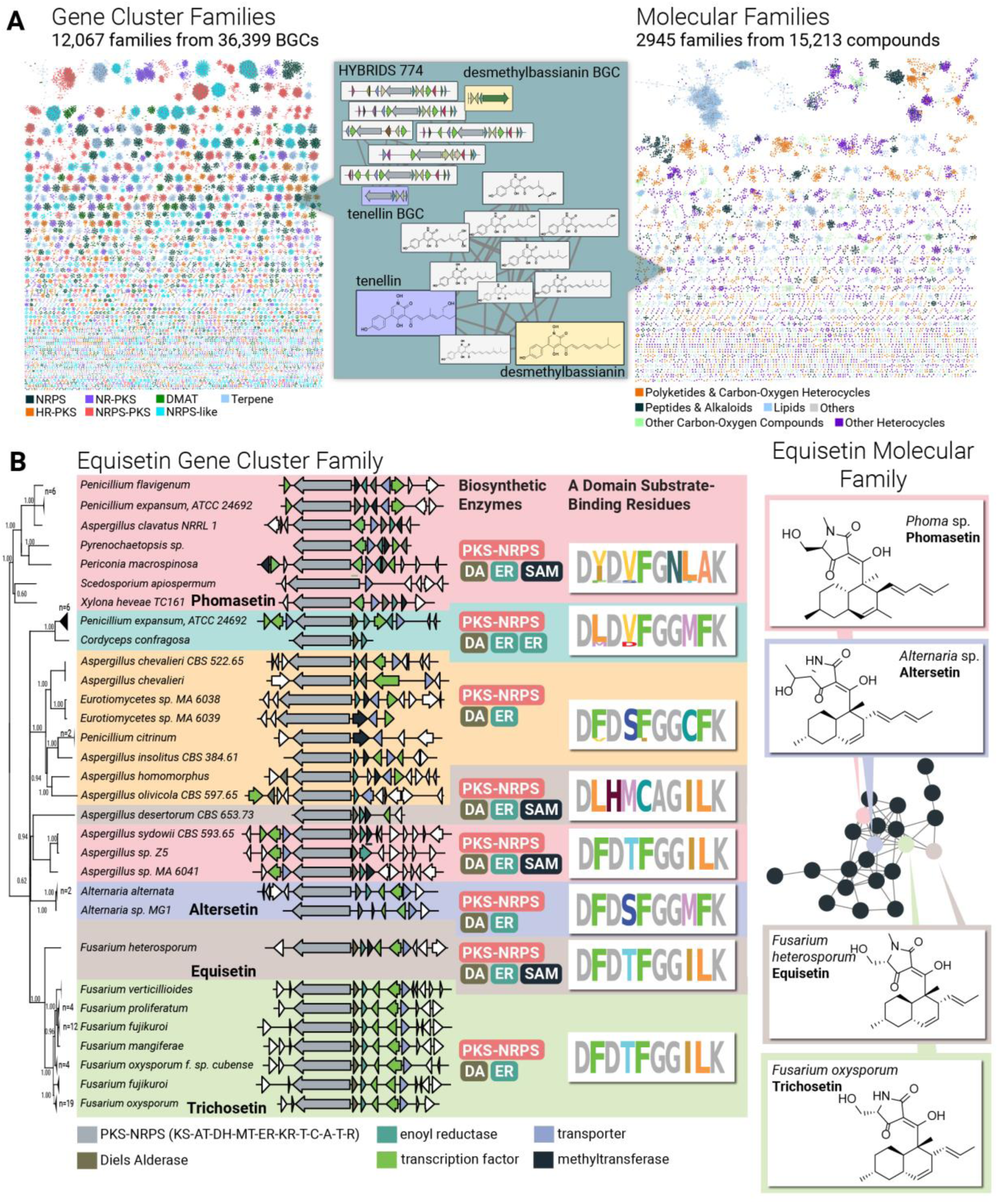
Large-scale analysis of fungal genome-encoded and known metabolite scaffolds. *(A)* Colliding large scale collections of fungal genetic content (at left) and fungal natural products (at right) using a network of gene cluster families (GCFs) interpreted from 1037 genomes (left) and 15,213 metabolites arranged into 2945 molecular families based on their Tanimoto similarity score (at right). Note that 92% of these 12,067 GCFs remain unassigned to their metabolite products. *(B)* Variations in adenylation domain substrate-binding residues and tailoring enzyme composition facilitate modifications to the equisetin GCF (left) and MF (right). The phylogram to the left represents a maximum likelihood tree based on the hybrid NRPS-PKS backbone enzyme. All branches in this tree have >50% bootstrap support.

We then sought to quantify the relationship between phylogeny and shared GCF content. To accomplish this, we used the protein sequence identity of 290 shared single-copy orthologous genes from the fungal BUSCO dataset (23) as a proxy for whole-genome distance; then we counted the fraction of GCFs shared within each genome in pairwise comparisons (**Fig. 2B**). A result was a clear relationship between genomic distance and shared GCF content, with an average of 75% shared GCFs at the species level, but less than 5% shared GCFs at taxonomic ranks higher than family (**Fig. 2C**). A similar trend exists for individual phyla and taxonomic classes (**Fig. S7**). Across the fungal kingdom, 76% of GCFs are species-specific and only 16% are genus-specific (**Fig. S8**), supporting the hypothesis that most BGCs enable fungi related at the species level to secure their respective ecological niches with highly specialized compounds (4).

### GCF-enabled annotation of fungal biosynthetic repertoire anchored by known BGCs

Identifying BGCs that have known metabolite products is an important component of genome mining, enabling researchers to prioritize known versus unknown biosynthetic pathways for discovery. These “genomic dereplication” efforts have been bolstered by the development of the MIBiG repository (24), which contains 213 fungal BGCs with known metabolites (at the time of these analyses, June 2019). When anchored with known BGCs, the GCF approach enables large-scale annotation of unstudied BGCs based on similarity to reference BGCs, identifying clusters likely to produce known metabolites or derivatives of knowns.

Within our dataset, 154 GCFs contained known BGCs from MIBiG, approximately 1% of the 12,067 total GCFs reported here (**Fig. S9**). These families collectively include a total of 2,026 BGCs (**Fig. S9**), an approximately 10-fold increase in the number of annotated BGCs over that available in MIBiG (24). To make this expanded set of annotated BGCs and their families available for routine genome mining, we created a section within the new *Prospect* website that highlights these newly annotated BGCs.

### Large-scale comparison of GCFs and fungal compounds

To assess the relationship between GCFs and their chemical repertoire, we next compared GCF-encoded scaffolds to a dataset of known fungal scaffolds. Analogous to our GCF analysis, we utilized network analysis of fungal metabolites, organizing these compounds into molecular families (MFs) based on Tanimoto similarity, a commonly used metric for determining chemical relatedness (25, 26). To directly relate GCF and MF-encoded metabolite scaffolds, we determined the relationship between chemical similarity and BGC similarity for a set of 154 fungal GCFs with known metabolite products (**Fig. S10**). We chose a MF similarity threshold that resulted in similar levels of chemical similarity represented by GCF and MF metabolite scaffolds.

Using this compound network analysis strategy, we organized a dataset of 15,213 fungal metabolites from the Natural Products Atlas (27) into 2,945 MFs (**Fig. 3A**). We annotated each compound within this network with chemical ontology information using ClassyFire, a tool for classifying compounds into a hierarchy of terms associated with structural groups, chemical moieties, and functional groups (see **Table S2** for a breakdown of this chemical ontology analysis for fungal metabolites) (28). The number of MF scaffolds (2,945) is only 25% the number of GCF-encoded scaffolds (12,067) in our 1000-genome dataset. This suggests that even this small genomic sampling of the entire fungal kingdom, estimated to have >1 million species (29), possesses biosynthetic potential that significantly dwarfs known fungal chemical space – not only in terms of individual *metabolites*, but also in terms of *metabolite scaffolds*. In this joint GCF-MF dataset, *molecular* families and *gene cluster* families represent complementary approaches for representing the same metabolite scaffold, such as the tenellin/desmethylbassianin structural class, whose GCF and MF contains both BGCs and compounds, respectively (**Fig. 3A**, middle). Exploring such pairings of GCF and MFs is a proven strategy for large-scale assignment of BGCs to their metabolite products (30), an activity that will provide a basis for improved compound discovery and for identifying the biosynthetic mechanisms fungi use for diversifying their bioactive scaffolds. An example of the latter is described below.

### Diversification of the equisetin scaffold inferred from gene cluster families

To further explore the link between metabolite scaffolds as represented by *molecular* and *gene cluster* families, we looked to the decalin-tetramic acids, a structural class well represented in our BGC and metabolite datasets. This structural class, including compounds such as equisetin, altersetin, phomasetin, and trichosetin (**Fig. S11**) (31-33), has a wide range of reported biological activities, including antibiotic, anti-cancer, phytotoxic, and HIV integrase inhibitory activity (34). We reasoned that further exploration of the decalin-tetramic acid structural class would yield insights into the biosynthetic mechanisms for variation of this bioactive scaffold by BGCs within the GCF.

Using *Prospect*, we identified two closely related GCFs (HYBRIDS_11/HYBRIDS_610) containing known BGCs responsible for biosynthesis of equisetin (35), trichosetin (36), and phomasetin (37) as well as BGCs from *Alternaria* likely responsible for the biosynthesis of altersetin found in multiple *Alternaria* species (32, 38). While most fungal GCFs are confined to single species or genera (**Fig. 2**), the equisetin GCF has an exceptionally broad phylogenetic distribution, with clusters found in the four Pezizomycotina classes Eurotiomycetes, Dothideomycetes, Xylonomycetes, and Sordariomycetes (**Fig. 3B**, left). The associated equisetin MF is likewise found in a variety of Dothideomycetes and Sordariomycetes (**Fig. 3B**, right).

The equisetin biosynthetic pathway involves three major steps: assembly of a decalin core via the action of polyketide synthase (PKS) enzyme domains and a Diels Alderase, formation of an amino acid-derived tetramic acid moiety catalyzed by NRPS domains, and N-methylation of the tetramic acid moiety (**Fig. S12**) (37, 39). While the domain structure of the PKS contained in the equisetin GCF remains consistent across fungi, differences in backbone enzyme amino acid sequence and the presence/absence of tailoring enzymes mediate structural variations to the scaffold. The PKS enzymes from *Fusarium oxysporum* and *Pyrenochaetopsis* sp. RK10-F058 share 50% sequence identity, which likely result in the additional ketide unit and C-methylation observed in equisetin vs. phomasetin (**Fig. 3B**). In the NRPS module of the hybrid NRPS-PKS, changes to adenylation domain substrate binding residues are predicted to mediate incorporation of serine (trichosetin, equisetin, and phomasetin) and threonine (altersetin). The *Aspergillus desertorum* BGC contains adenylation domain substrate binding residues that are highly variant from those found in other clusters within the GCF, indicating its tetramic acid moiety is likely diversified with a different amino acid. The equisetin GCF contains additional variations in the number of enoyl reductase enzymes (one additional in the uncharacterized *Penicillium expansum* clade), indicating possible differences to degree of saturation, and a methyltransferase that is expected to mediate changes in tetramic acid N-methylation.

This pattern of biosynthetic variation within a GCF resulting in metabolite diversification suggests that exploring such pairs of GCFs and MFs with knowledge of their taxonomic distribution will be valuable to guide genome mining in the identification of new analogs of compounds with proven therapeutic or agrochemical value. The equisetin GCF is one of only 90 GCFs (representing 0.75% of total GCFs) within our dataset that spanned multiple taxonomic classes (**Table S3**). This includes bioactive scaffolds such PR-toxin, swainsonine, chaetoglobosin, and cytochalasin (**Fig. S13**) which contain variations in tailoring enzyme composition expected to diversify these scaffolds. Given the observed biosynthetic diversity within such “multi-class” GCFs, exploring such pairs of GCFs and MFs represents an attractive approach for discovering new analogs of bioactive metabolites.

### A “bird’s eye” view of fungal versus bacterial biosynthetic space

Having surveyed GCFs across the fungal kingdom, we sought to compare and contrast this genomic and chemical repertoire to the well-established bacterial canon. We gathered 5,453 bacterial genomes whose BGCs were publicly available in the antiSMASH bacterial BGCs database (40), resulting in a dataset of 24,024 bacterial BGCs to compare to our dataset of 36,399 fungal BGCs. To visualize the biosynthetic space encompassed by these BGCs, we determined the frequency of protein domains within BGCs for each major taxonomic group (see **SI Methods**). Principle Component Analysis (PCA) of these encoded BGCs showed a phylogenetic bias in this biosynthetic space, with bacteria and fungi occupying distinct regions (**Fig. 4A**).

**Figure 4.**
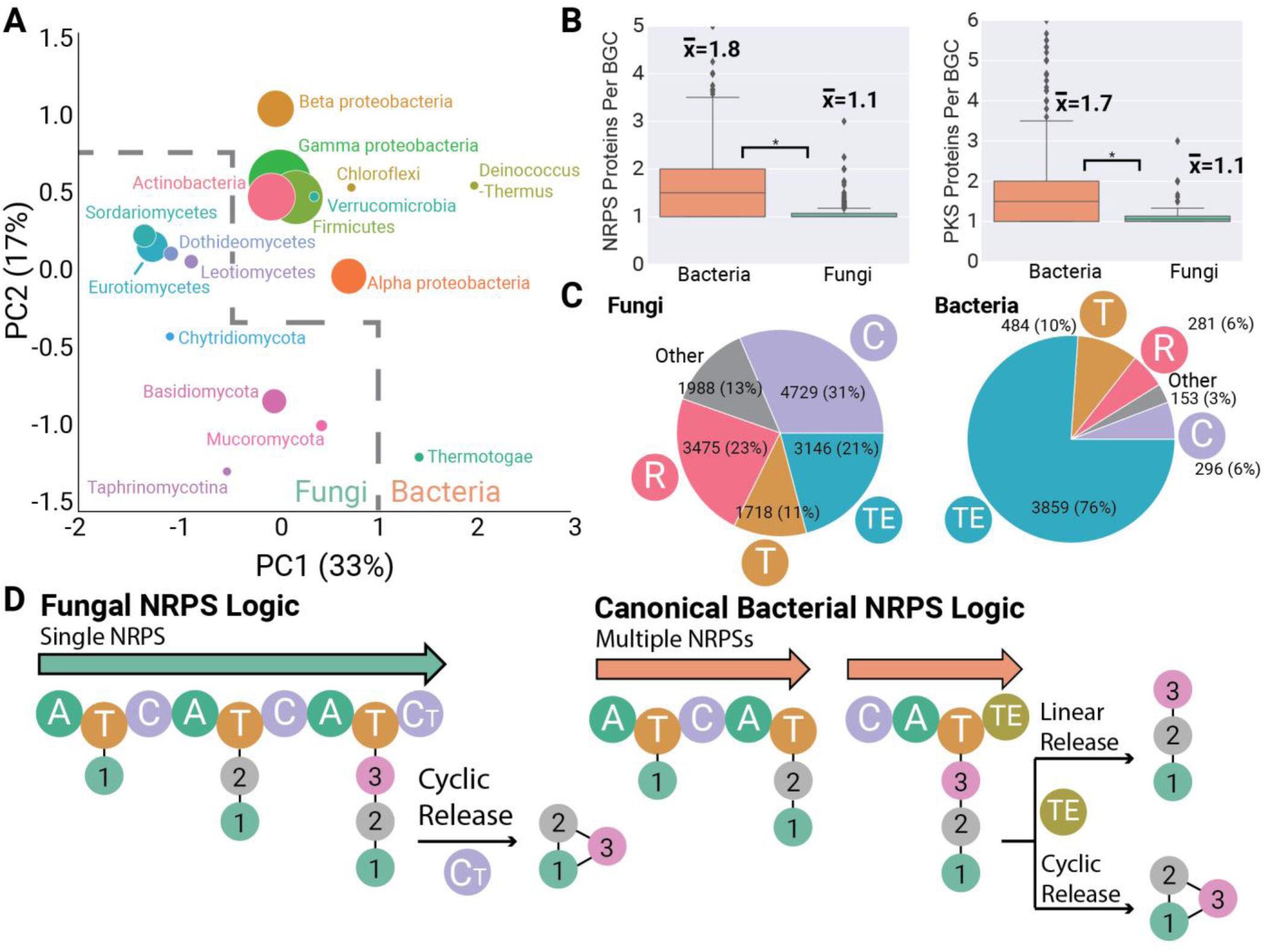
Fungal biosynthetic gene clusters are distinct from their canonical bacterial counterparts. *(A)* Principle Component Analysis (PCA) of 36,399 fungal and 24,024 bacteria biosynthetic gene clusters (BGCs), with points sized according to the number of BGCs analyzed. Fungal and bacterial taxonomic groups occupy distinct regions of this biosynthetic space. *(B)* Fungal and bacterial BGCs differ in backbone enzyme composition, with fungal NRPS and PKS clusters typically encoding only a single backbone, compared to multiple backbone enzymes found in bacterial BGCs. *(C)* Fungal and bacterial NRPS BGCs differ dramatically in their use of termination domains for release of peptide intermediates. *(D)* Fungal NRPS logic is distinct from bacterial canon. Most fungal NRPS pathways involve a single NRPS enzyme that utilizes a terminal condensation domain to produce a cyclic peptide. In contrast, bacterial NRPS enzymes contain multiple NRPS enzymes that operate in a colinear fashion and typically utilize thioesterase domains to produce linear or cyclic peptides.

We observed dramatic differences in bacterial versus fungal NRPS and PKS assembly line logic. Consistent with prior studies of iterative fungal PKS enzymes (41), fungal PKS BGCs typically encode a single backbone PKS enzyme, while bacterial PKS BGCs contain a median of 1.7 PKS backbone enzymes per cluster (**Fig. 4B**, right). Surprisingly, fungal NRPS BGCs also usually encode a single backbone enzyme, compared to multiple backbone enzymes more typically observed in bacterial systems (**Fig. 4B**, left). Fungal NRPS and PKS enzymes also average ∼150% the size of bacterial backbones (**Fig. S14**). In addition to these contrasting backbone enzyme compositions, we observed systematic differences in the top NRPS domain organizations (**Fig. S15**), particularly in NRPS termination domains (**Fig. 4C**). The most common fungal NRPS termination domains are C-terminal condensation domains, recently found to catalyze release of peptide intermediates via intramolecular cyclization (42-44). The next most common are terminal thioester reductase domains that perform either reductive release to aldehydes or alcohols or release via cyclization (45). This is in stark contrast to bacterial NRPS BGCs, which most commonly terminate with type I thioesterase domains that release intermediates as linear or cyclic peptides (**Fig. 4C**).

These collective differences between fungal and bacterial BGCs show systematic differences in NRPS biosynthetic logic between these two kingdoms. In bacterial NRPS canon, a pathway is comprised of multiple NRPS genes whose chromosomal order (and the order of catalytic domain “modules” within the encoded polypeptide) corresponds to the order of amino acid monomers in the metabolite product (**Fig. 4D**, right) (46). In the field of bacterial natural products, the use of this “collinearity rule” to predict metabolite scaffolds is commonplace (19, 47, 48); however, the large number of exceptions to this rule reduces the accuracy of these predictions. The prototypical fungal NRPS (**Fig. 4D** (**Fig. 4D**) primarily involves the action of biosynthetic domains within the same backbone enzyme, rather than multiple NRPS backbones acting in concert. This suggests that future efforts to predict fungal NRPS scaffolds will be able to largely bypass the need to account for permutations of multiple NRPS genes, raising the possibility of increased predictive performance compared to bacteria.

### Uncovering distinct natural product reservoirs

Having shown that fungi and bacteria are distinct biosynthetically, we sought to compare these genomics-based insights to the chemical space of known metabolites. We added 9,382 bacterial compounds to our dataset of 15,213 fungal metabolites, analyzing these bacterial compounds using the same network analysis and chemical ontology workflow described above. We performed PCA to visualize the chemical space of major fungal and bacterial taxonomic groups within this compound dataset (further described in **SI Methods**).

PCA of bacterial and fungal compounds (**Fig. 5A**) revealed a trend that parallels our analysis of fungal and bacterial biosynthetic space (**Fig. 4A**). Bacteria and fungi occupy separate regions of chemical space, differing dramatically in terms of chemical ontology superclass, a high-level descriptor of general structural type (**Fig. 5B**). Fungi have twice the frequency of lipids and nearly twice the frequency of heterocyclic compounds, a structural group that includes aromatic polyketide-related moieties such as furans and pyrans. Many of the chemical moieties and structural classes that are highly enriched in bacteria or fungi are vital in bioactive scaffolds. This includes moieties such as the bacterial aminoglycoside antibiotics (49), thiazoles present in the bacterial anti-cancer bleomycin family (50), and the steroid ring that forms the core scaffold of steroid drugs such as the fungal metabolite fusidic acid (51) (**Fig. 5B**). PCA loadings plots similarly reveal differences between bacterial and fungal chemical space, including a high prevalence of peptide-associated chemical ontology terms in bacteria, and lipid and aromatic polyketide terms in fungi (**Fig. S16**).

**Figure 5.**
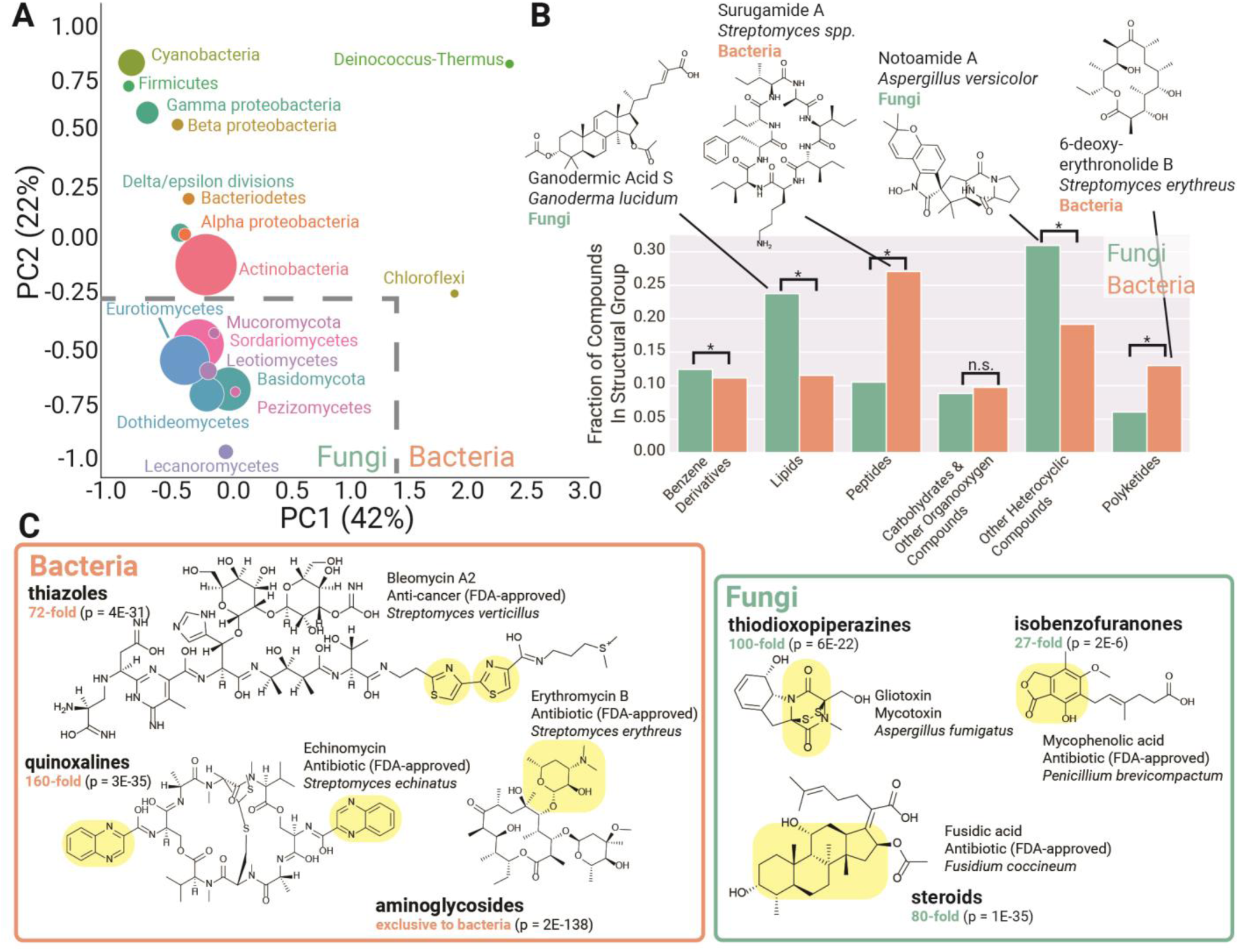
Bacteria and fungi are distinct sources for natural product scaffolds. *(A)* Principal Component Analysis (PCA) of 24,595 known bacterial and fungal compounds, with points sized according to the number of compounds. Fungal and bacterial taxonomic groups occupy distinct regions in this representation of chemical space for natural products. *(B)* Quantitative comparison of structural classifications in bacterial vs fungal compounds. *(C)* Bacteria and fungi represent distinct pools for bioactive compounds and scaffolds. Selected chemical moieties enriched and characteristic of each taxonomic group are highlighted in yellow. The fold enrichment of the chemical moiety is indicated in green, with p-values from a Chi-Squared test indicated.

Within the fungal kingdom, we observed differences in PCA of the chemical repertoire of major taxonomic groups (**Fig. S17**). Pezizomycotina classes grouped together in chemical space, largely due to a higher proportion of polyketide and peptide-related chemical moieties (**Fig. S18**). Basidiomycota are distinct chemically, possessing a much higher proportion of chemical moieties and descriptors associated with terpenes and other lipids. These observations based on chemical space are consistent with the higher proportion of NRPS and PKS BGCs within Pezizomycotina and the prevalence of terpene BGCs within Basidiomycota groups such as Agaricomycotina (**Fig. 2B**), and further supported by PCA of fungal BGCs, in which fungal phyla represent distinct groups (**Figs. S19** and **S20**).

## Discussion

### A framework for exploring fungal scaffolds using gene cluster families

The GCF approach enables the systematic mapping of the biosynthetic repertoire encoded by large groups of fungal genomes. The fungal kingdom is a wealth of untapped biosynthetic potential, with the 1000 genomes analyzed here representing a reservoir of >12,000 new GCF-encoded scaffolds. This genome dataset is only a small subset of the >1 million predicted fungal species (29), indicating that the total biosynthetic potential of the fungal kingdom far surpasses that assembled here.

By organizing biosynthetically related BGCs into families, the GCF approach provides a means of cataloguing and dereplicating genome-encoded MFs. In the field of bacterial natural products discovery, this GCF paradigm has been expanded for automated linking of GCFs to MFs detected by metabolomics and molecular networking analysis, enabling high-throughput genome mining from industrial-scale strain collections (5, 7, 29, 52). Establishing the GCF approach for fungal genomes lays the groundwork for similar GCF-driven large-scale compound discovery efforts from fungi.

### Data-driven prospecting for fungal natural products

Large-scale genome sequencing projects such as the 1000 Fungal Genomes project, whose stated goal is sampling every taxonomic family within Fungi (53), will uncover a large amount of biosynthetic and chemical novelty. However, as 76% of fungal GCFs are species- and 16% are genus-specific, such genome sequencing efforts focused on taxonomic families will miss the majority of GCFs. Additional large-scale efforts to sample this biosynthetic space based on “depth” rather than “breadth” is suggested to more efficiently access these genomes. Future “1000-genomes” projects, now feasible for academic research groups due to ever-decreasing genome sequencing costs, should focus on expanding this dataset with species-level sequencing of taxonomic groups.

The GCF approach provides a means of selecting fungi for compound and BGC discovery via approaches such as heterologous expression (54) based not on taxonomic or phylogenetic markers, but with a strategy that focuses on efficient sampling of biosynthetic pathways. The distribution of GCFs shows groups of organisms with shared GCFs (**Fig. S6**), and sampling based on these organism “groups” reduces the number of genomes required to capture the majority of fungal biosynthetic space. Our simulated sampling based on shared GCFs indicated that 80% of GCFs from the 386 Eurotiomycete genomes are represented in a sample of only 145 genomes. By contrast, to represent the same number of GCFs, species-level sampling required 189 genomes and random sampling required 263 genomes (**Fig. S21**). This indicates that the GCF approach can be used as a roadmap for systematic characterization of new fungal biosynthetic pathways and their compounds.

### Unearthing new medicines

These analyses of both chemical and biosynthetic space show that bacteria and fungi represent chemically distinct sources for natural products discovery. Interestingly, fungal compounds are closer to FDA-approved compounds than bacterial compounds in terms of several chemical properties, including three out of four “Lipinsky Rule of Five” properties often used as guidelines for predicting oral bioavailability (**Fig. S22**) (55). While many of the most successful natural products violate these rules of thumb, these data suggest that fungal metabolites may be more “druglike” than those occupying bacterial chemical space.

Compound discovery efforts should be initiated with the understanding that different biological sources will yield distinct chemical space and different types of metabolite scaffolds. The fungal kingdom is rich in aromatic polyketides, while bacteria harbor a higher proportion of peptidic scaffolds. Within the fungal kingdom, Basidiomycota is a rich reservoir of terpene scaffolds, while BGC-rich Pezizomycotina classes are a richer source of polyketides and peptides. These data suggest that distinct taxonomic groups not only possess the capacity for different metabolite scaffolds, but also different *types* of scaffolds.

## Conclusion

We have mapped the landscape of 12,067 GCFs across 1037 fungal genomes, revealing the phylogenetic distribution of these families and establishing a framework for high-throughput genome mining from fungi. This framework introduces a new fundamental biosynthetic unit – the gene cluster family – for cataloguing and annotating the rapidly increasing number of fungal genomes available. Network analysis at the GCF level advances the field of fungal genome mining with an approach scalable to industrial-scale strain collections, providing a new approach for systematically mapping known and unknown fungal biosynthetic space and associated metabolite scaffolds. The GCF paradigm further provides an atlas for exploring metabolite scaffolds and their derivatives, enabling targeted genome mining focused on scaffolds with proven value. These collective analyses reveal that genomes across the fungal kingdom represent a rich resource for discovery of new natural products. In both under-explored and well-studied fungal taxa, a wide variety of metabolite scaffolds awaits discovery, and the ever-decreasing cost of genome sequencing will help usher in a wave of large-scale fungal genome mining efforts that rival those currently underway in bacteria.

## Acknowledgments

This work was supported by the National Center for Complementary and Integrative Health (R01 AT009143). L.K.C. was supported by the National Institute of General Medical Sciences (F32 GM134679). M.T.D. was supported by U.S. Department of Agriculture, National Institute of Food and Agriculture (USDA NIFA) postdoctoral fellowship award 2019-67012-29662.

