## Supplemental Information for "An Interpreted Atlas of Biosynthetic Gene Clusters from 1000 Fungal Genomes"

### An Interpreted Atlas of Biosynthetic Gene Clusters and Their Families from 1000 Fungal Genomes.

#### Supplementary Methods

*Genome dataset*

A set of 1,037 fungal genome assemblies was downloaded from GenBank and the JGI Genome Portal. The subphylum *Saccharomycotina* was excluded, as a large portion of genomes in this taxonomic group are known to lack biosynthetic gene clusters relevant to this study. Genomes were downloaded 11/15/2018. All genomes were processed using the command-line version of antiSMASH 4 with the fungal setting and otherwise default parameters (1). This resulted in 36,399 predicted biosynthetic gene clusters, to which we added 213 quality fungal gene cluster sequences with known metabolite products from the MIBiG database (2).

*Gene cluster family network creation*

We utilized a workflow similar to that utilized by the recently-developed tool BiG-SCAPE (3). Predicted biosynthetic gene clusters (GenBank files from antiSMASH) were converted to arrays of predicted protein domains using the HMMER suite tool *hmmsearch* using the Pfam 31.0 database. This database of protein domain Hidden Markov Models was supplemented with models corresponding to fungal polyketide product template domains (TIGR04532) and terpene cyclase domain (NCBI CDD cd00687) based on aligned sequences present in NCBI (4).

The Jaccard similarity between all pairs of gene cluster domain arrays was calculated using the Python package scikit-learn, and any pairs with Jaccard similarity less than 0.1 were excluded from further analysis (3, 5). Gene cluster pairs that passed this Jaccard similarity threshold were compared based on sequence identity. Gene clusters were first separated into biosynthetic type based on the presence or absence of protein domains as defined in **Table S4**. This resulted in gene clusters classified as nonribosomal peptide synthase (NRPS), highly-reducing polyketide synthase (HR-PKS), nonreducing polyketide synthase (NR-PKS), hybrid NRPS-PKS, NRPS-like, dimethylallyl transferase (DMAT), or terpene. We chose to use mutually inclusive classifications (e.g. both NRPS and DMAT for a single cluster), which enabled downstream analysis in cases where the antiSMASH-predicted boundaries encompassed multiple gene clusters in close chromosomal proximity. We determined sequence identity between all pairs of gene clusters within a given biosynthetic class by performing sequence-profile alignments of backbone enzyme using the HMMER suite tool *hmmalign* (4). For a given pair of gene clusters, sequence identity was calculated as the mean sequence identity between pairs of backbone enzyme domains. In the case of gene clusters with more than one of a given backbone enzyme domain, the Hungarian matching algorithm was used identify the configuration of domain pairs with the highest sequence similarity, and only these domain pairs were used (3).

A final sequence similarity score was calculated as:

$$Similarity\ Score = \sqrt{0.8 * sequence\ identity + 0.2 * Jaccard\ similarity}.$$

DBSCAN clustering was used on the resulting distance matrix, with an epsilon value of 0.50. This workflow thus resulted in a network or graph structure, in which nodes represent gene clusters and subgraphs represent gene cluster families. All data was output to a PostgreSQL database to interface with the web portal. Comparisons of BGC similarity to compound similarity in **Fig. S10** were performed using structures associated with 213 known fungal BGCs from the MIBiG database (2). Structures were converted to chemical fingerprints using the Morgan Fingerprint method with a radius of 2 and compared via Tanimoto similarity as implemented in RDKit (6).

*Web portal creation (Prospect: prospect-fungi.com)*

The developed web portal uses the Python web framework Django as a REST API backend and the TypeScript framework Angular as a frontend interface. The main access points to the site include a table of gene cluster families (GCFs) and a table of genomes analyzed in this study. Each GCF page, accessible via a unique GCF ID (e.g. NRPS_42), shows gene clusters within the family, along with corresponding taxonomy information. When a gene cluster is selected, a panel shows various metadata corresponding to the cluster and a Cytoscape network for visualizing the GCF. A tab within the panel for Gene Info shows metadata for a selected gene, along with predicted protein domains. Individual protein domains can be selected to view a description of the domain from the Pfam website. Sequences corresponding to the gene cluster, all proteins, and predicted domains are available for download from the site. An additional Organisms page enables similar access to GCFs through specific genomes of interest.

*Phylogenetic trees*

All species trees were based on the BUSCO fungal dataset (7). The HMMER suite tool hmmsearch was used to extract protein sequences corresponding to BUSCO Hidden Markov Models (HMMs).(4) Extracted sequences were aligned using MAFFT with the –auto parameter.(8) Positions with >10% gaps were removed from the alignments using TrimAL (9). Trimmed alignments were concatenated, resulting in single aligned sequences for each genome. Phylogenetic trees were constructed in MEGA using Neighbor Joining, 500 bootstrap iterations, and the James-Taylor-Thornton model (10).

*Cheminformatics Analyses*

A database of known metabolites and their biological sources was downloaded from the Natural Products Atlas on December 10, 2020 (11). Compounds were classified by taxonomy using each metabolite record’s genus and species information by referencing the NCBI taxonomy database, downloaded October 22, 2019. Predicted chemical ontology information was determined from each compound's InChi representation using the ClassyFire REST API (12). Compounds were converted to chemical fingerprints using the Morgan Fingerprint method with a radius of 2, as implemented in RDKit (6). The Tanimoto similarity function is RDKit was used to calculate the pairwise distance between compounds (6). A cutoff of 0.6 Tanimoto similarity was used to create compound families within the network. Bacterial and fungal metabolites were compared to FDA-approved compounds from the DrugBank database (13). All chemical descriptors and properties were calculated using RDKit (6).

*Principal Component Analysis (PCA)*

Principal Component Analyses were conducted on two datasets and data subsets thereof. To evaluate differences in the frequencies of biosynthetic domains across fungi, each fungal gene cluster was converted into an array representing the frequency of 14,350 protein domains across each taxonomic group. Taxonomic groups with less than ten genomes were removed from analysis to avoid skewing frequency calculations. Differences in fungal and bacterial chemical space were evaluated using the frequencies of chemical ontology descriptors in molecules downloaded from the aforementioned Natural Products Atlas chemical database (11), including only taxonomic groups with 5 or more compounds in the downloaded database. Final chemometric analyses were conducted using Sirius version 10.0 (Pattern Recognition Systems AS).

| Taxon | NRPS | HYBRID | HRPKS | TERPENE | NRPSLIKE | NRPKS | DMAT | Genomes analyzed | Per-genome average |
|---|---|---|---|---|---|---|---|---|---|
| **Pucciniomycotina** | 2.0 | 0.0 | 0.0 | 0.6 | 0.6 | 0.0 | 0.0 | 25 | 3.2 |
| **Ustilaginomycotina** | 3.0 | 0.6 | 0.1 | 0.0 | 3.6 | 0.9 | 0.3 | 32 | 8.5 |
| **Agaricomycotina** | 1.6 | 0.6 | 0.1 | 6.1 | 4.5 | 1.0 | 0.2 | 173 | 14.1 |
| **Pezizomycotina** | 9.0 | 5.4 | 4.7 | 4.5 | 7.9 | 7.1 | 1.2 | 721 | 39.8 |
| **Taphrinomycotina** | 1.2 | 0.0 | 0.0 | 0.5 | 1.1 | 0.0 | 0.0 | 12 | 2.8 |
| **Mucoromycota** | 1.1 | 0.2 | 0.0 | 1.6 | 2.6 | 0.0 | 0.0 | 36 | 5.5 |
| **Zoopagomycota** | 3.8 | 0.1 | 0.1 | 0.3 | 0.7 | 0.1 | 0.0 | 16 | 5.1 |
| **Blastocladiomycota** | 1.5 | 0.0 | 0.0 | 0.0 | 0.5 | 0.0 | 0.0 | 2 | 2 |
| **Chytridiomycota** | 9.1 | 0.9 | 0.8 | 0.1 | 1.6 | 1.2 | 0.0 | 12 | 13.7 |
| **Microsporidia** | 0.0 | 0.0 | 0.0 | 0.0 | 0.0 | 0.0 | 0.0 | 2 | 0 |
| **Cryptomycota** | 0.0 | 0.0 | 0.0 | 0.0 | 0.0 | 1.0 | 0.0 | 1 | 1.0 |

**Table S1.** Genomes analyzed in this study and the distribution of their gene clusters classified by biosynthetic type.

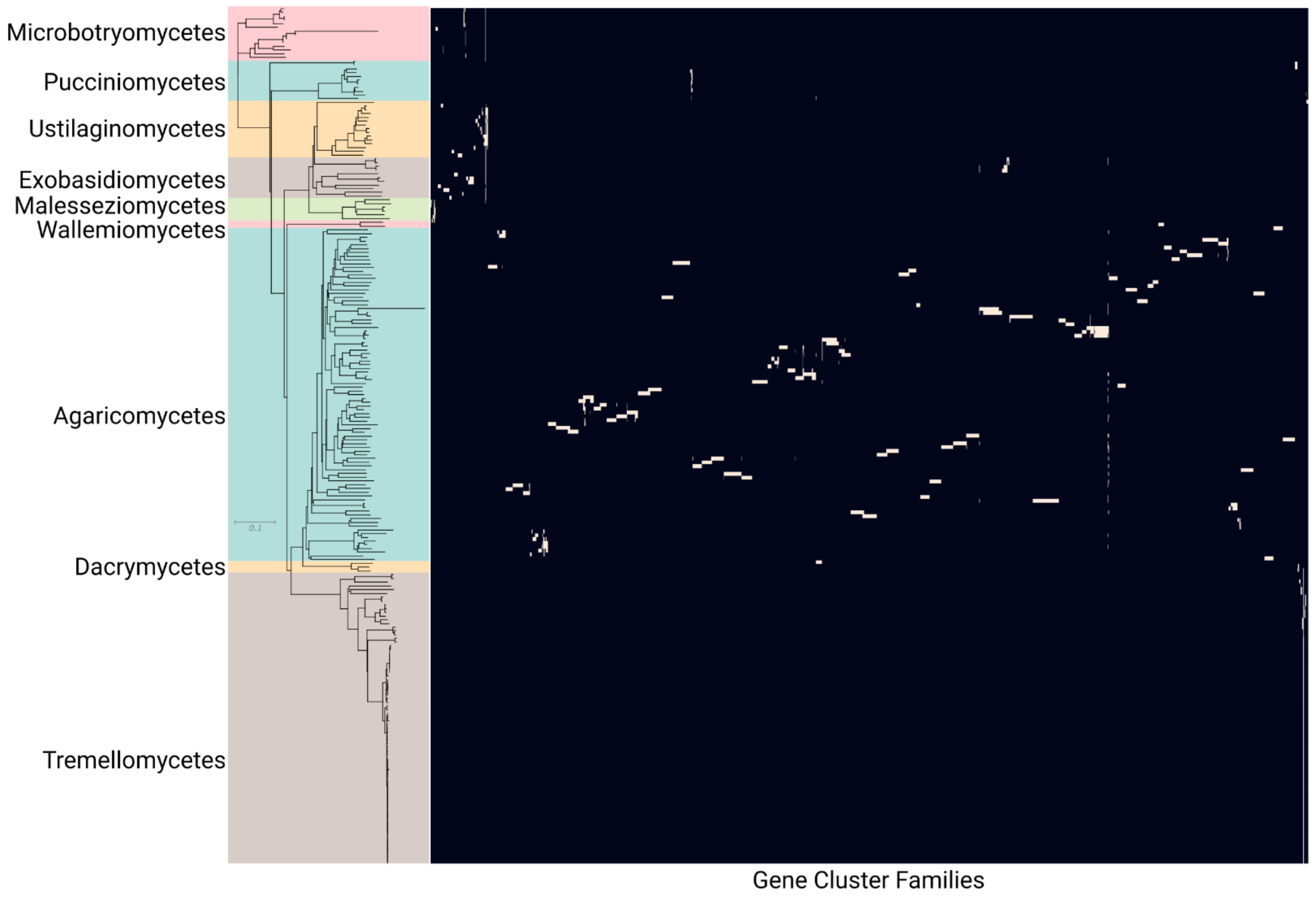


**Fig. S1.** Distribution of 1933 gene cluster families (GCFs) across Basidiomycota. The phylogram to the left shows a Neighbor Joining tree based on 290 orthologous genes, with branches with less than 50% bootstrap support collapsed. Genomes are colored by class, according to NCBI taxonomy information. Genomes within Tremellomycetes are largely composed of subspecies of *Cryptococcus neoformans* and *Cryptococcus gatti* and show little variation in GCF content. Within other classes of Basidiomycota, the majority of GCFs are species- or genus-specific. Several GCFs are distributed across entire classes or shared by organisms within different classes.

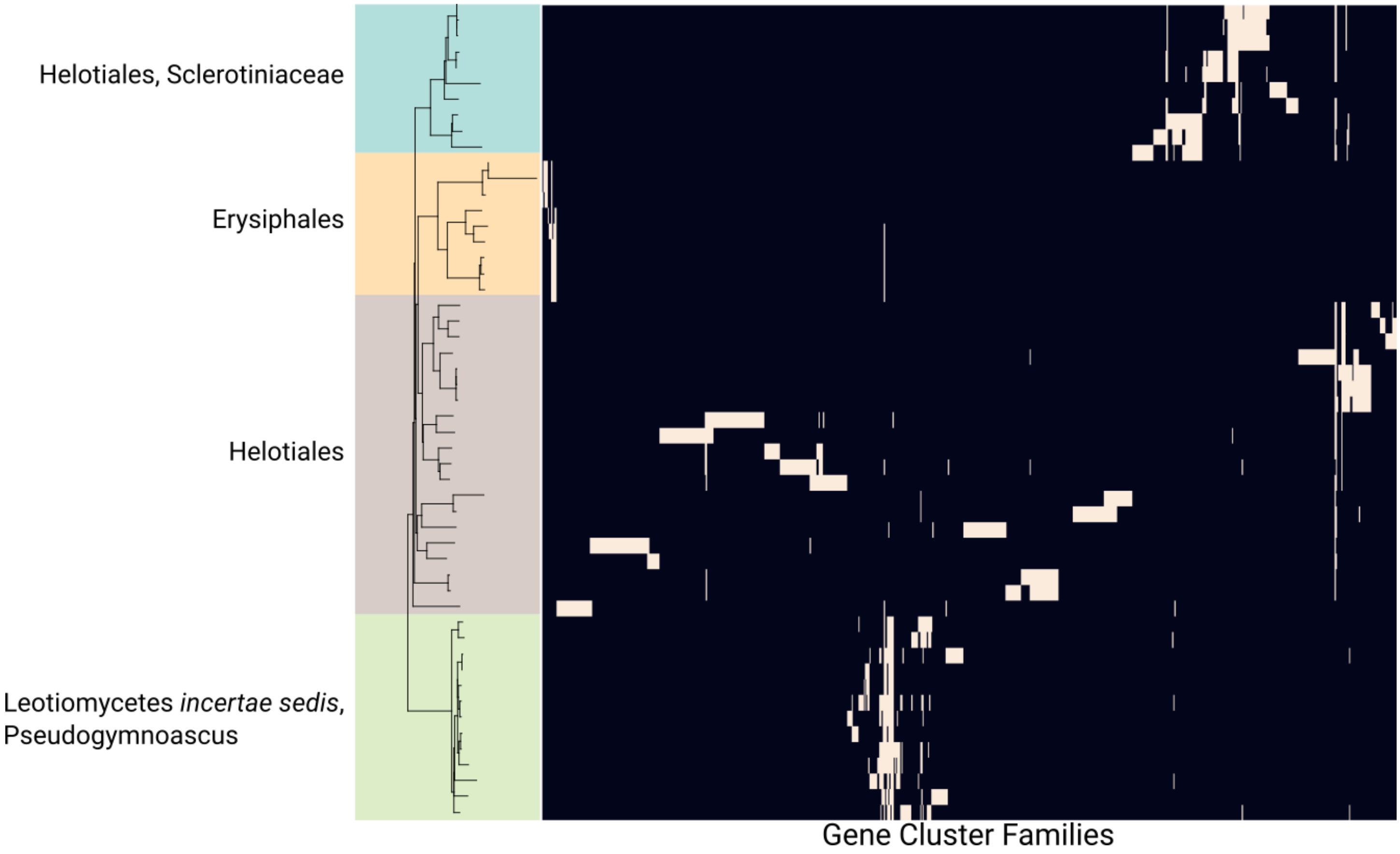


**Fig. S2.** Distribution of 822 gene cluster families (GCFs) across Leotiomycetes. The phylogram to the left shows a Neighbor Joining tree based on 290 orthologous genes, with branches with less than 50% bootstrap support collapsed. Genomes are colored by taxonomic order, according to NCBI taxonomy information.

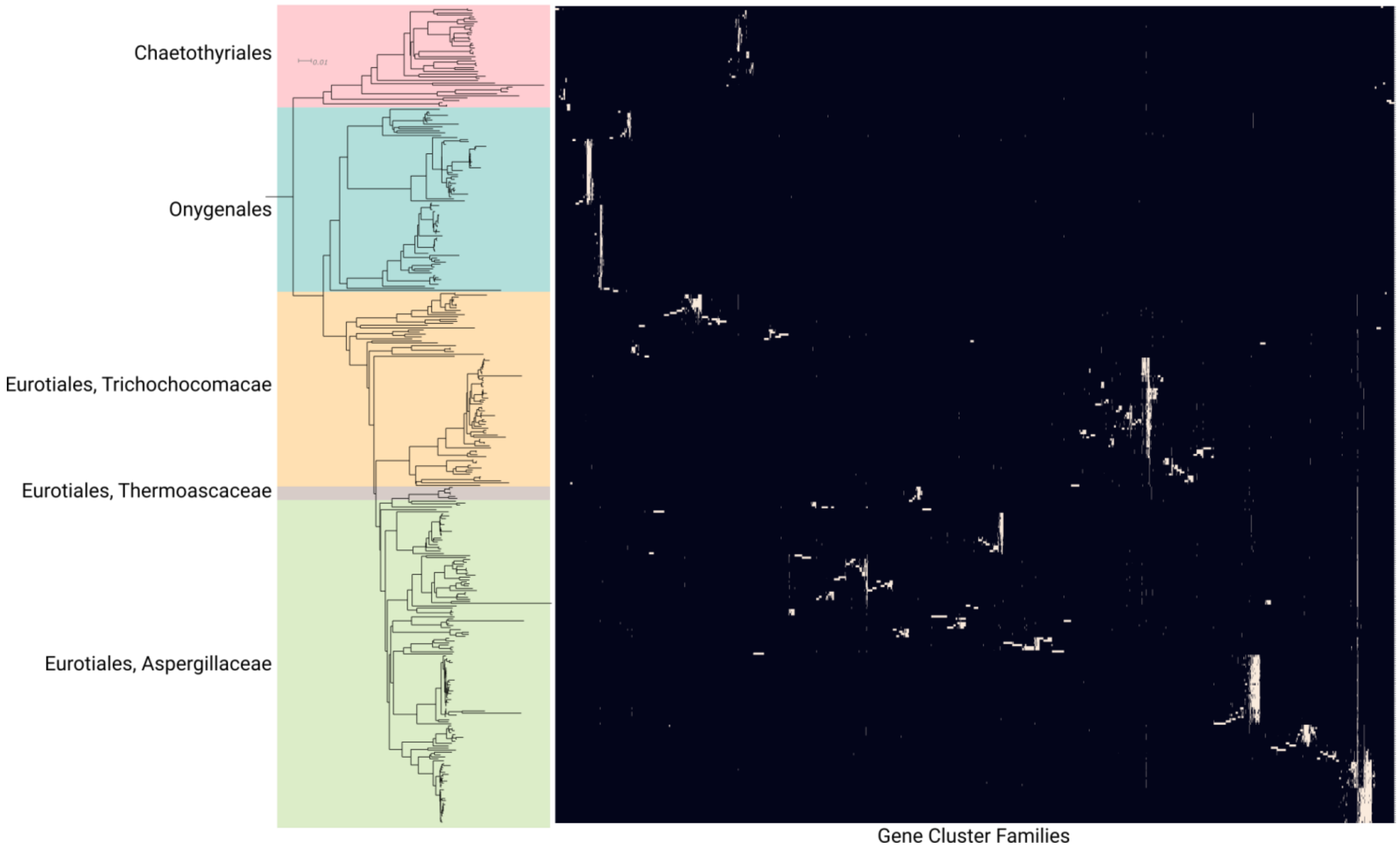


**Fig. S3.** Distribution of 4926 gene cluster families (GCFs) across Eurotiomycetes. The phylogram to the left shows a Neighbor Joining tree based on 290 orthologous genes, with branches with less than 50% bootstrap support collapsed. Genomes are colored by taxonomic order, according to NCBI taxonomy information.

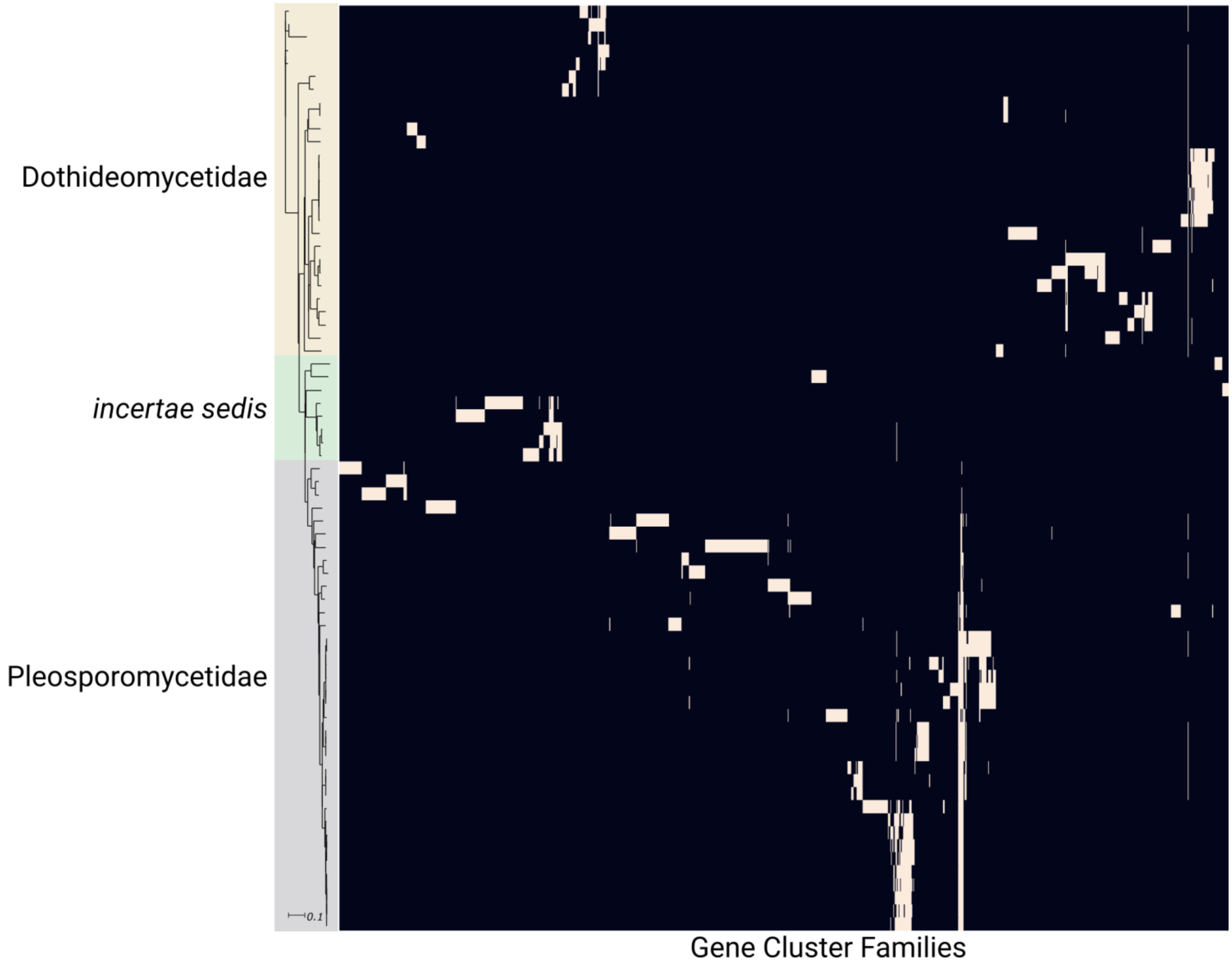


**Fig. S4.** Distribution of 1176 gene cluster families (GCFs) across Dothideomycetes. The phylogram to the left shows a Neighbor Joining tree based on 290 orthologous genes, with branches with less than 50% bootstrap support collapsed. Genomes are colored by taxonomic order, according to NCBI taxonomy information.

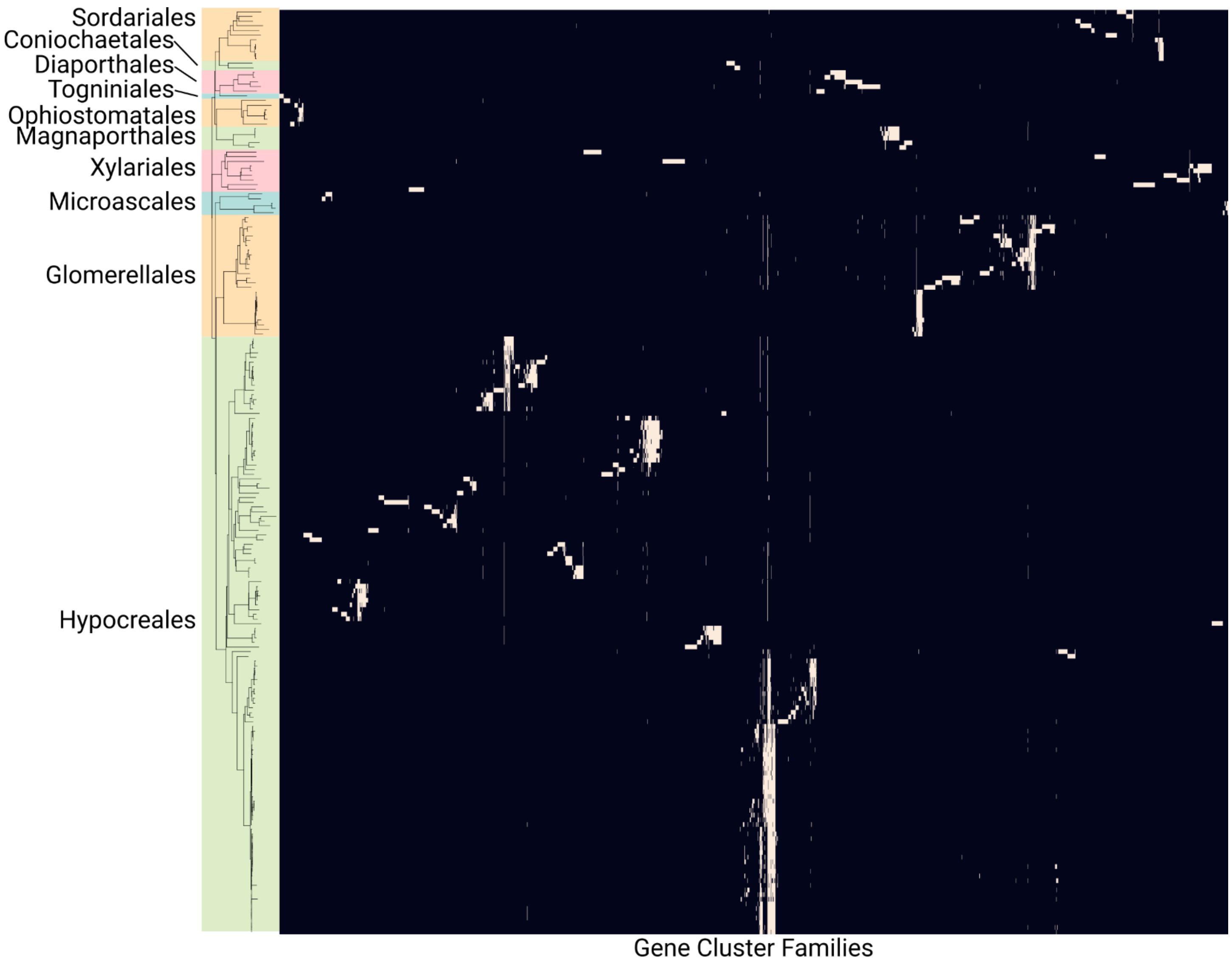


**Fig. S5.** Distribution of 2884 gene cluster families (GCFs) across Sordariomycetes. The phylogram to the left shows a Neighbor Joining tree based on 290 orthologous genes, with branches with less than 50% bootstrap support collapsed. Genomes are colored by taxonomic order, according to NCBI taxonomy information.

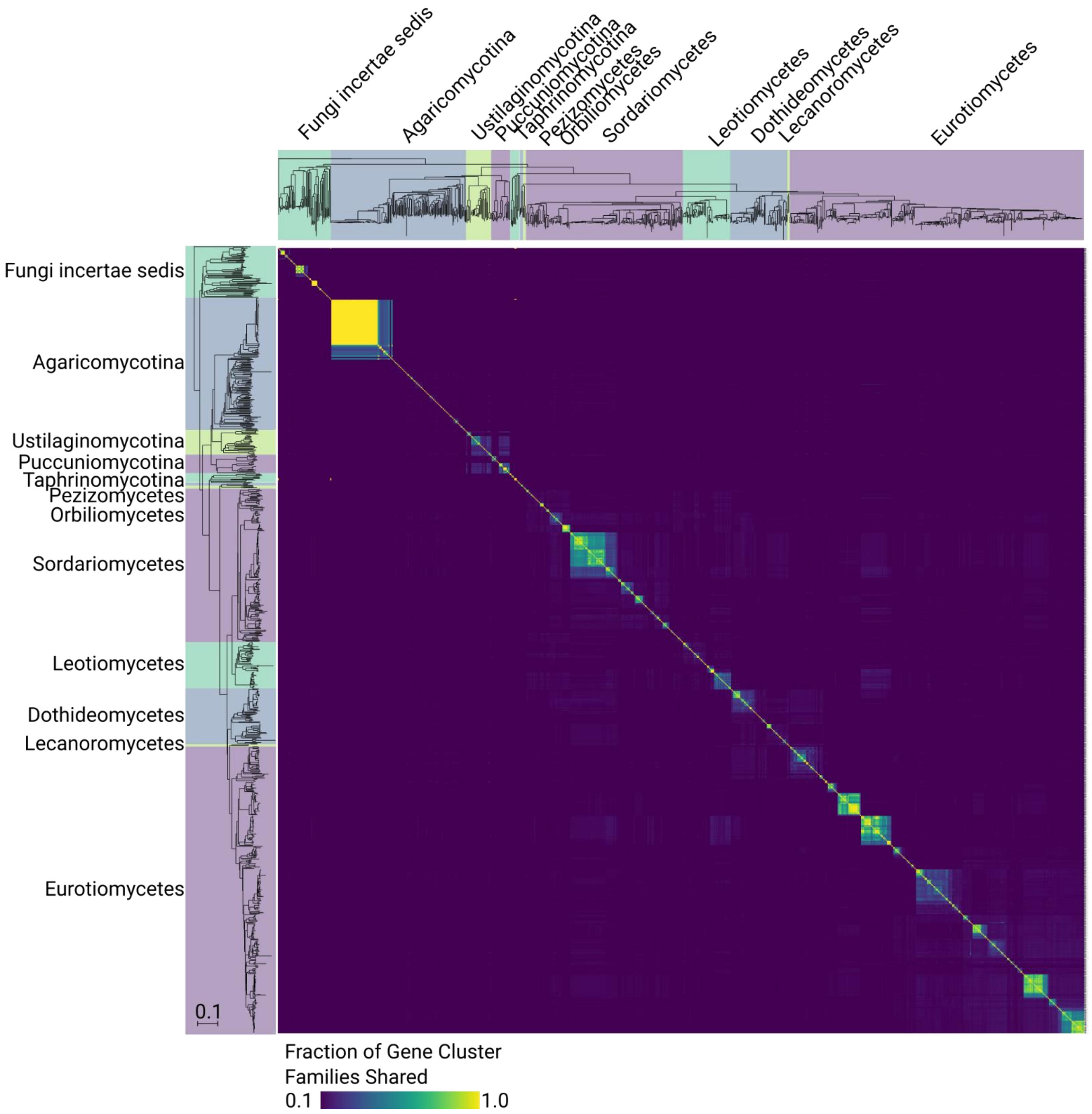


**Fig. S6.** Relationship between phylogeny and shared gene cluster family (GCF) content. The phylogram to the left shows a Neighbor Joining tree based on 290 orthologous genes, with branches with less than 50% bootstrap support collapsed. Genomes within Pezizomycotina are labeled by taxonomic class, according to NCBI taxonomy information. Other genomes are labeled by subphylum, according to NCBI taxonomy information.

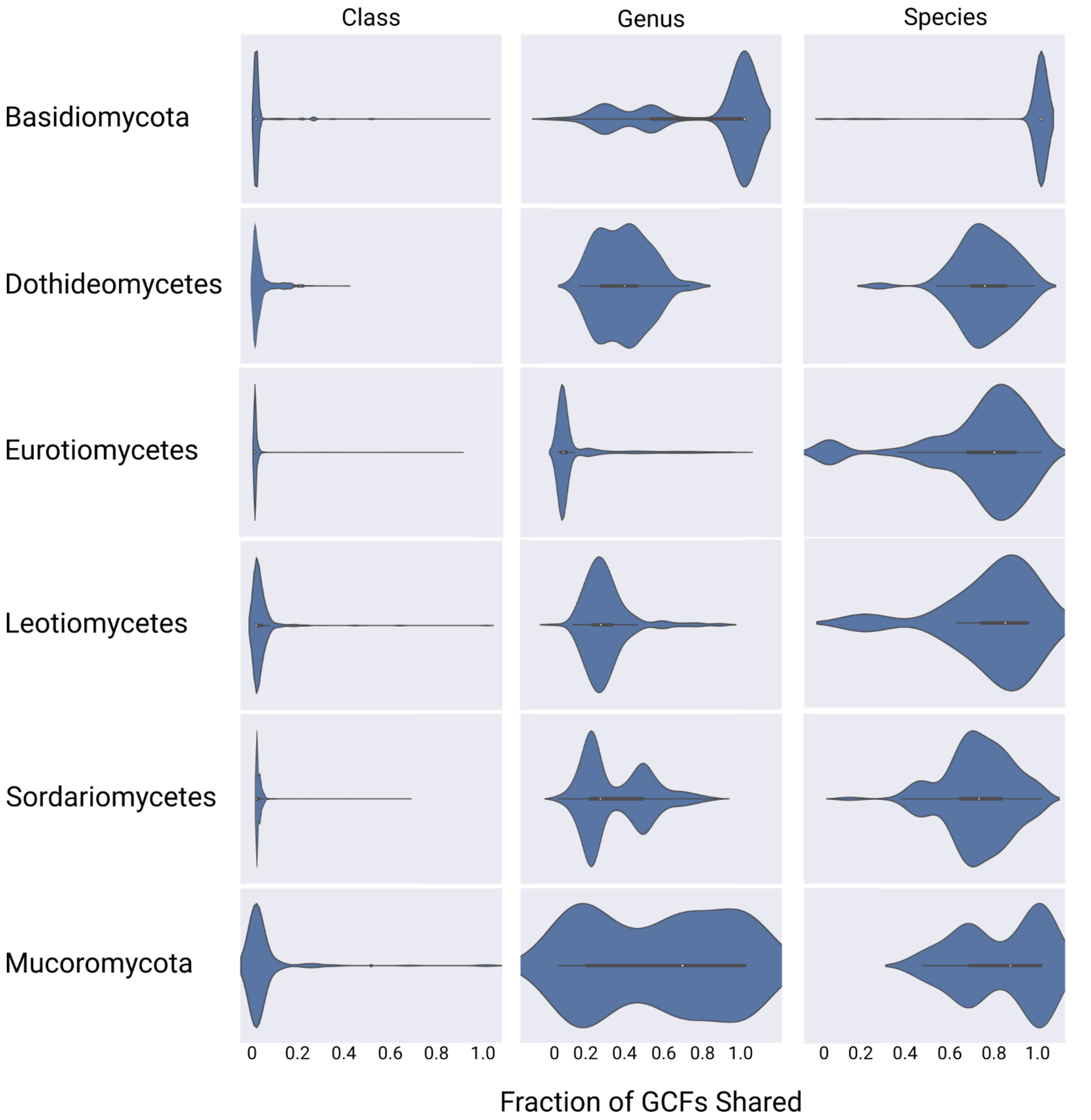


**Fig. S7**. Relationship between phylogeny and GCFs in six major taxonomic groups. The violin plots represent the fraction of gene cluster families (GCFs) shared by pairs of genomes within the given taxonomic groups. Each genome pair was given a mutually-exclusive classification of same-species, same-genus, or same-class, and the fraction of GCFs shared for each genome pair was determined.

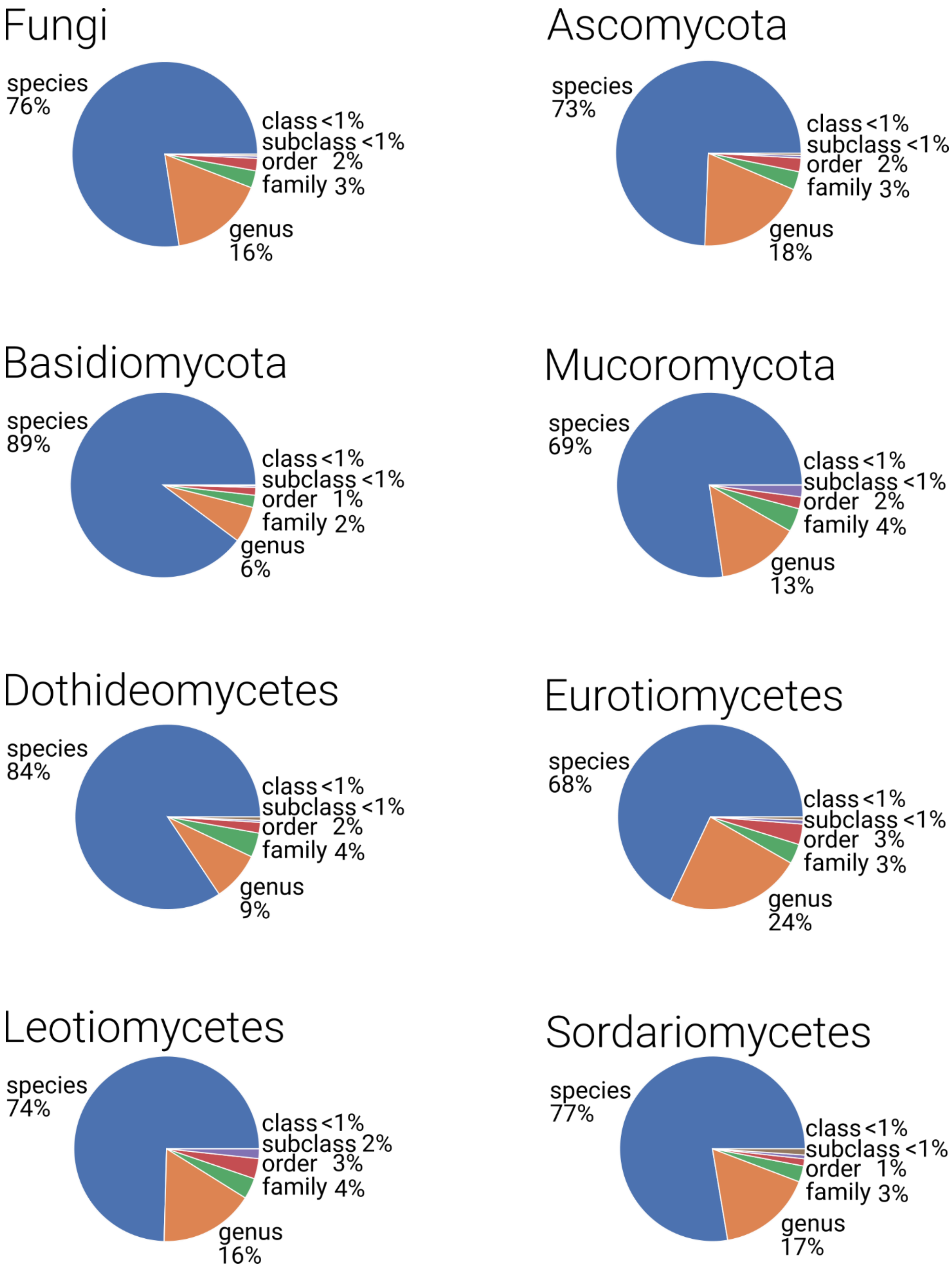


**Fig. S8**. Fungal gene cluster families (GCFs) are largely species-specific. Each GCF within the given taxonomic group was classified based on highest taxonomic rank shared by organisms with the GCF (i.e. species-specific, genus-specific, family-specific, etc.). Depending on taxonomic group, GCFs are between 68-89% species-specific.

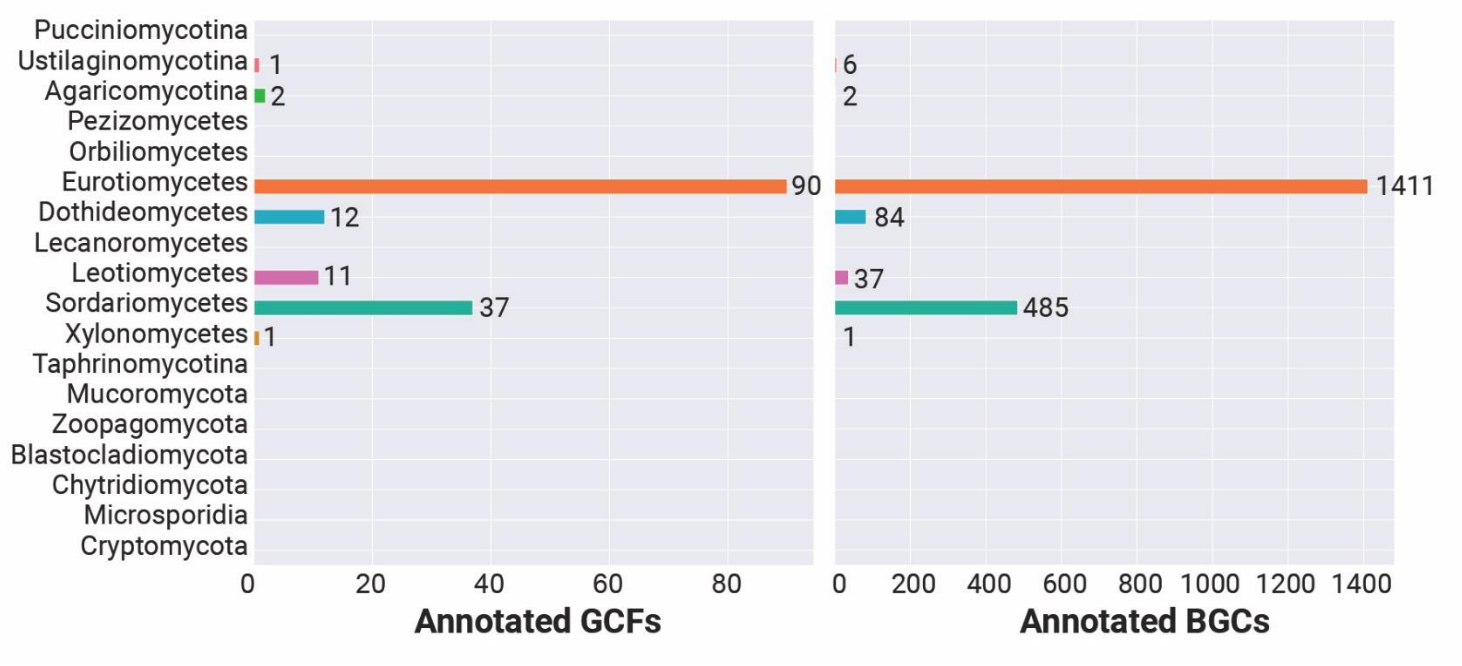


**Fig. S9**. Using the GCF approach for automated annotation of fungal BGCs with putative metabolite scaffolds. Across the taxonomic groups examined, a total of 154 GCFs contain reference BGCs with known metabolite products. At the level of individual clusters, these amounts to 2,026 BGCs annotated based on their presence in GCFs with known metabolite scaffolds.

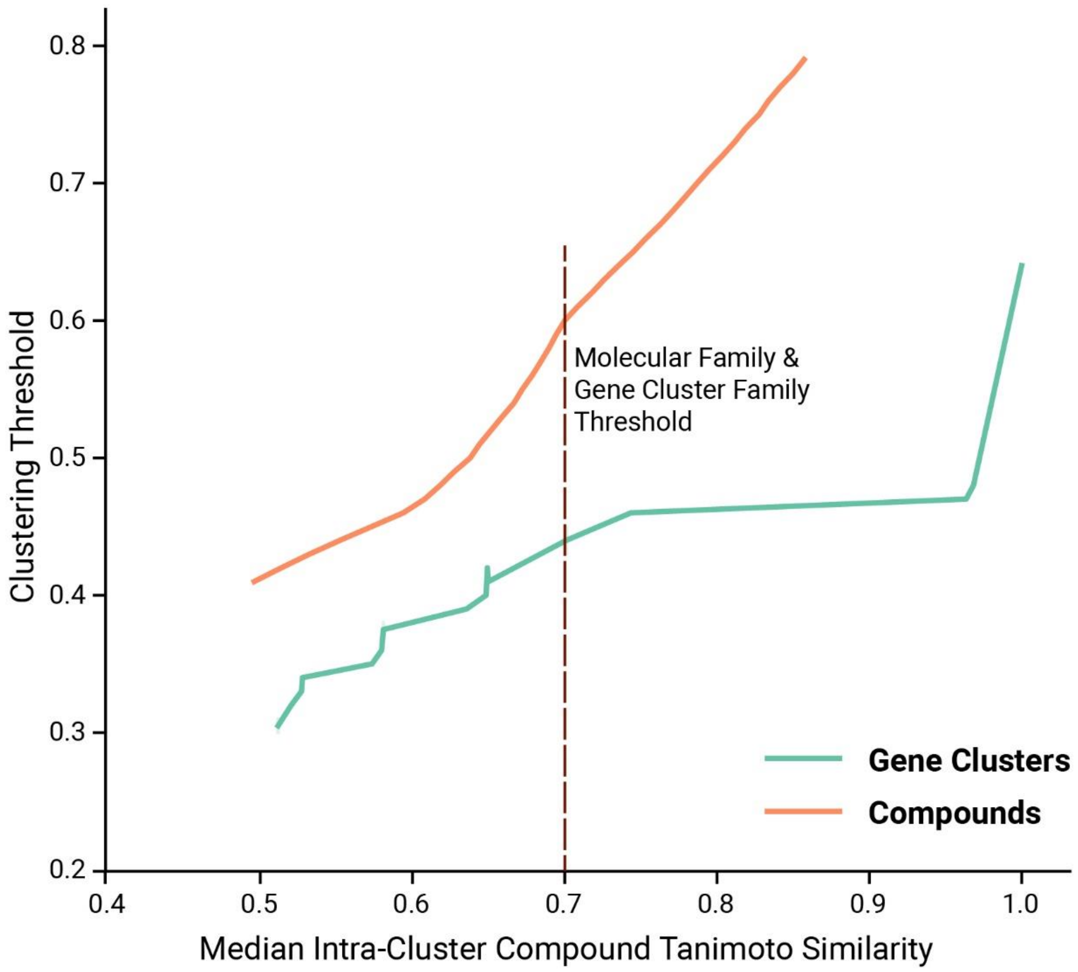


**Fig. S10.** Comparison of metabolite scaffold chemical space covered by molecular families (MFs) and gene cluster families (GCFs). At each clustering threshold, the median Tanimoto similarity of known compounds within GCFs and MFs was determined. A median intra-cluster Tanimoto similarity of 0.7 was chosen, corresponding to GCF and MF similarity thresholds of 0.45 and 0.6, respectively.

**Table. S2**. Chemical ontology-based classification of metabolites from *Aspergillus fumigatus*. Each chemical ontology entry in the table contains the major ontology superclass in bold, followed by other chemical ontology terms.

| Metabolite Name | Structure | Chemical Ontology Terms |
|---|---|---|
| 1,2-dihydro-16-O-deacetylhelvolic acid 21,16-lactone | 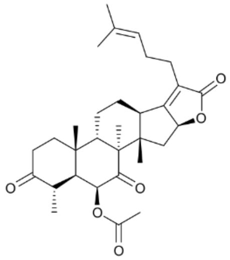 | **Lipids and lipid-like molecules**, Steroids and steroid derivatives, Steroid lactones, Steroid esters, 7-oxosteroids, 3-oxo-5-alpha-steroids, Alpha-acyloxy ketones, Dicarboxylic acids and derivatives, Butenolides, Enoate esters, Lactones, Cyclic ketones, Oxacyclic compounds, Organic oxides, Hydrocarbon derivatives |
| 11-methyl-11-hydroxyldodecanoic acid amide | 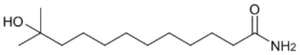 | **Organic oxygen compounds**, Organooxygen compounds, Alcohols and polyols, Tertiary alcohols, Carboximidic acids, Organopnictogen compounds, Organonitrogen compounds, Hydrocarbon derivatives |
| 11-O-methylpseurotin A | 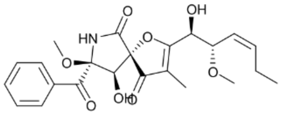 | **Organic oxygen compounds**, Organooxygen compounds, Carbonyl compounds, Ketones, Aryl ketones, Phenylketones, Alkyl-phenylketones, Benzoyl derivatives, Aryl alkyl ketones, Pyrrolidine-2-ones, Furanones, Vinylogous esters, Secondary carboxylic acid amides, Secondary alcohols, Lactams, Oxacyclic compounds, Dialkyl ethers, Azacyclic compounds, Organonitrogen compounds, Organic oxides, Hydrocarbon derivatives |
| 13-oxoverruculogen | 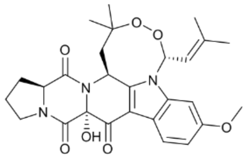 | **Organoheterocyclic compounds**, Indoles and derivatives, Pyridoindoles, Beta carbolines, Alpha amino acids and derivatives, Indoles, 2,5-dioxopiperazines, Aryl alkyl ketones, Anisoles, N-alkylpiperazines, Alkyl aryl ethers, Vinylogous amides, Tertiary carboxylic acid amides, Pyrrolidines, Pyrroles, Heteroaromatic compounds, Lactams, Dialkyl peroxides, Oxacyclic compounds, Azacyclic compounds, Alkanolamines, Hydrocarbon derivatives |
| 2,4,6,8-Decatetraenedioic acid, mono[2-methoxy-4-methylene-3-[2-methyl-3-(3-methyl-2-butenyl)-2-oxiranyl]cyclohex-1-yl] ester | 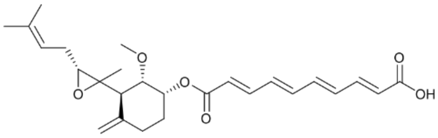 | **Lipids and lipid-like molecules**, Fatty Acyls, Fatty acids and conjugates, Medium-chain fatty acids, Fatty acid esters, Epoxy fatty acids, Branched fatty acids, Unsaturated fatty acids, Dicarboxylic acids and derivatives, Enoate esters, Oxacyclic compounds, Epoxides, Dialkyl ethers, Carboxylic acids, Organic oxides, Hydrocarbon derivatives, Carbonyl compounds |
| 2-chloro-1, 3, 8-trihydroxy-6-methylanthrone | 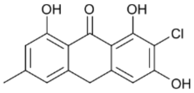 | **Benzenoids**, Anthracenes, Halophenols, Aryl ketones, 1-hydroxy-4-unsubstituted benzenoids, 1-hydroxy-2-unsubstituted benzenoids, Aryl chlorides, Vinylogous acids, Polyols, Organochlorides, Organic oxides, Hydrocarbon derivatives |
| 3-hydroxy-2,5-toluquinone | 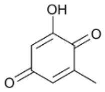 | **Organic oxygen compounds**, Organooxygen compounds, Carbonyl compounds, Ketones, Cyclic ketones, Quinones, Benzoquinones, P-benzoquinones, Vinylogous acids, Enols, Organic oxides, Hydrocarbon derivatives |
| 4,8,10,14-tetramethyl-6-acetoxy-14-[16-acetoxy-19-(20,21-dimethyl)-18-ene]phenanthrene-1-ene-3,7-dione | 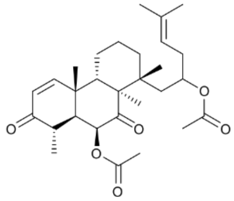 | **Benzenoids**, Phenanthrenes and derivatives, Hydrophenanthrenes, Cyclohexenones, Alpha-acyloxy ketones, Dicarboxylic acids and derivatives, Carboxylic acid esters, Organic oxides, Hydrocarbon derivatives |

| Metabolite Name | Structure | Chemical Ontology Terms |
| --- | --- | --- |
| 5-N-acetylardeemin | 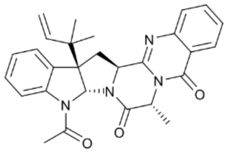 | **Organoheterocyclic compounds**, Indoles and derivatives, Pyrroloindoles, Quinazolines, Indoles, Pyrimidones, Benzenoids, Acetamides, Tertiary carboxylic acid amides, Heteroaromatic compounds, Pyrrolidines, Pyrroles, Lactams, Azacyclic compounds, Carbonyl compounds, Hydrocarbon derivatives, Organic oxides, Organonitrogen compounds, Organopnictogen compounds |
| 6-hydroxytryprostatin | 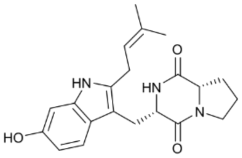 | **Organic acids and derivatives**, Carboxylic acids and derivatives, Amino acids, peptides, and analogues, Amino acids and derivatives, Alpha amino acids and derivatives, 3-alkylindoles, Hydroxyindoles, 2,5-dioxopiperazines, 1-hydroxy-2-unsubstituted benzenoids, N-alkylpiperazines, Substituted pyrroles, Tertiary carboxylic acid amides, Pyrrolidines, Heteroaromatic compounds, Secondary carboxylic acid amides, Lactams, Azacyclic compounds, Carbonyl compounds, Hydrocarbon derivatives, Organic oxides, Organonitrogen compounds, Organopnictogen compounds |
| Asperfumigatin | 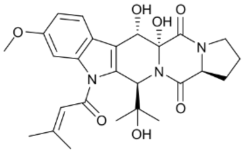 | **Organoheterocyclic compounds**, Indoles and derivatives, Pyridoindoles, Beta carbolines, Alpha amino acids and derivatives, 3-alkylindoles, Anisoles, 2,5-dioxopiperazines, N-alkylpiperazines, Alkyl aryl ethers, Substituted pyrroles, Tertiary carboxylic acid amides, Tertiary alcohols, Pyrrolidines, Heteroaromatic compounds, Secondary alcohols, Lactams, Azacyclic compounds, Alkanolamines, Organopnictogen compounds, Organic oxides, Hydrocarbon derivatives, Carbonyl compounds |
| Cephalimysin A | 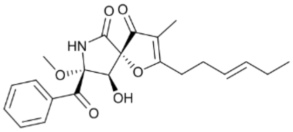 | **Organic oxygen compounds**, Organooxygen compounds, Carbonyl compounds, Ketones, Aryl ketones, Phenylketones, Alkyl-phenylketones, Benzoyl derivatives, Aryl alkyl ketones, Pyrrolidine-2-ones, Furanones, Vinylogous esters, Secondary carboxylic acid amides, Secondary alcohols, Lactams, Oxacyclic compounds, Azacyclic compounds, Organonitrogen compounds, Organic oxides, Hydrocarbon derivatives |
| Fumagiringillin | 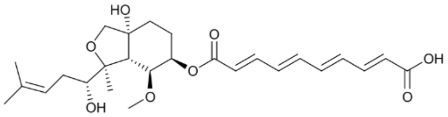 | **Organoheterocyclic compounds**, Isobenzofurans, Medium-chain fatty acids, Branched fatty acids, Hydroxy fatty acids, Heterocyclic fatty acids, Fatty acid esters, Unsaturated fatty acids, Dicarboxylic acids and derivatives, Tetrahydrofurans, Tertiary alcohols, Enoate esters, Secondary alcohols, Cyclic alcohols and derivatives, Oxacyclic compounds, Carboxylic acids, Dialkyl ethers, Organic oxides, Carbonyl compounds, Hydrocarbon derivatives |
| Fumiformamide | 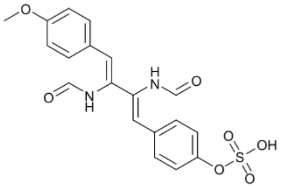 | **Lignans**, neolignans and related compounds, Phenylsulfates, Phenoxy compounds, Methoxybenzenes, Anisoles, Alkyl aryl ethers, Sulfuric acid monoesters, Propargyl-type 1,3-dipolar Carboximidic acids, Organonitrogen compounds, Organic oxides, Hydrocarbon derivatives |
| Fumifungin | 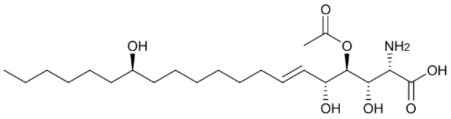 | **Lipids and lipid-like molecules**, Fatty Acyls, Fatty acids and conjugates, Long-chain fatty acids, L-alpha-amino acids, Hydroxy fatty acids, Beta hydroxy acids and derivatives, Amino fatty acids, Unsaturated fatty acids, Dicarboxylic acids and derivatives, Secondary alcohols, Carboxylic acid esters, Amino acids, Polyols, Carboxylic acids, Organopnictogen compounds, Organic oxides, Monoalkylamines, Hydrocarbon derivatives, Carbonyl compounds |
| Fumigaclavine C | 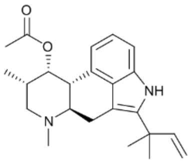 | **Alkaloids and derivatives**, Ergoline and derivatives, Clavines and derivatives, Indoloquinolines, Benzoquinolines, Pyrroloquinolines, 3-alkylindoles, Isoindoles and derivatives, Aralkylamines, Substituted pyrroles, Piperidines, Benzenoids, Heteroaromatic compounds, Trialkylamines, Amino acids and derivatives, Carboxylic acid esters, Monocarboxylic acids and derivatives, Azacyclic compounds, Carbonyl compounds, Hydrocarbon derivatives, Organic oxides, Organopnictogen compounds |

| Metabolite Name | Structure | Chemical Ontology Terms |
|---|---|---|
| Fumigatonin | 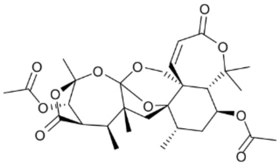 | **Lipids and lipid-like molecules**, Prenol lipids, Sesquiterpenoids, Abscisic acids and derivatives, Terpene lactones, Tetracarboxylic acids and derivatives, Oxepanes, Ortho esters, Ketals, Carboxylic acid orthoesters, Gamma butyrolactones, 1,3-dioxanes, Tetrahydrofurans, Enoate esters, Oxacyclic compounds, Organic oxides, Hydrocarbon derivatives, Carbonyl compounds |
| Fumigatoside B | 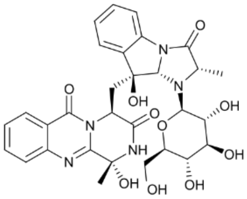 | **Organic oxygen compounds**, Organooxygen compounds, Carbohydrates and carbohydrate conjugat Glycosyl compounds, Glycosylamines, Hexoses, Quinazolines, Alpha amino acids and derivatives, Indoles and derivatives, Pyrimidones, Oxanes, Imidazolidinones, Benzenoids, Tertiary carboxylic acid amides, Tertiary alcohols, Heteroaromatic compounds, Cyclic carboximidic acid Secondary alcohols, Lactams, Hemiaminals, Propargyl-type 1,3-dipolar Polyols, Oxacyclic compounds, Azacyclic compounds, Primary alcohols, Organopnictogen compounds, Organic oxides, Hydrocarbon derivatives, Carbonyl compounds |
| Fumiquinazoline A | 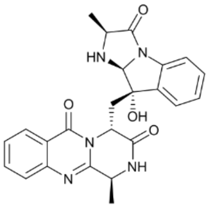 | **Organoheterocyclic compounds**, Diazanaphthalenes, Benzodiazines, Quinazolines, Alpha amino acids and derivatives, Indoles and derivatives, Pyrimidones, Imidazolidinones, Benzenoids, Tertiary carboxylic acid amides, Tertiary alcohols, Heteroaromatic compounds, Lactams, Secondary carboxylic acid amides, Dialkylamines, Azacyclic compounds, Organopnictogen compounds, Organic oxides, Hydrocarbon derivatives, Carbonyl compounds |
| Fumiquinone A | 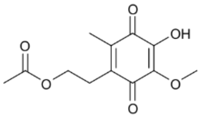 | **Lipids and lipid-like molecules**, Prenol lipids, Quinone and hydroquinone lipids, Prenylquinones, Ubiquinones, P-benzoquinones, Vinylogous esters, Vinylogous acids, Carboxylic acid esters, Monocarboxylic acids and derivatives, Organic oxides, Hydrocarbon derivatives |
| Fumisoquin A | 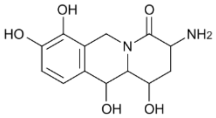 | **Organoheterocyclic compounds**, Tetrahydroisoquinolines, Alpha amino acids and derivatives, Piperidinones, Delta lactams, Aralkylamines, Aminopiperidines, 1-hydroxy-4-unsubstituted benzenoid 1-hydroxy-2-unsubstituted benzenoids, Tertiary carboxylic acid amides, Secondary alcohols, Polyols, Azacyclic compounds, Organopnictogen compounds, Organic oxides, Monoalkylamines, Hydrocarbon derivatives, Carbonyl compounds |
| Fumitremorgin A | 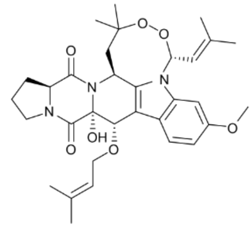 | **Organoheterocyclic compounds**, Indoles and derivatives, Pyridoindoles, Beta carbolines, 3-alkylindol Alpha amino acids and derivatives, 2,5-dioxopiperazines, Anisoles, Alkyl aryl ethers, N-alkylpiperazines, Heteroaromatic compounds, Pyrroles, Tertiary carboxylic acid amides, Pyrrolidines, Lactams, Dialkyl peroxides, Dialkyl ethers, Azacyclic compounds, Oxacyclic compounds, Alkanolamines, Hydrocarbon derivatives, Carbonyl compounds, Organopnictogen compounds |
| GERI-BP001 | 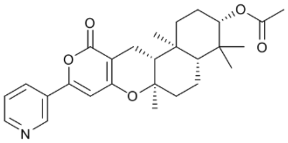 | **Organoheterocyclic compounds**, Naphthopyrans, Naphthalenes, Alkyl aryl ethers, Pyranones and d Pyridines and derivatives, Vinylogous esters, Heteroaromatic compounds, Lactones, Carboxylic acid est Oxacyclic compounds, Monocarboxylic acids and derivatives, Azacyclic compounds, Organic oxides, Hydrocarbon derivatives, Carbonyl compounds, Organonitrogen compounds, Organopnictogen compo |

| Metabolite Name | Structure | Chemical Ontology Terms |
|---|---|---|
| Gliotoxin G | 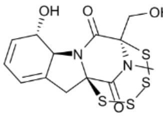 | **Organic acids and derivatives**, Carboxylic acids and derivatives, Amino acids, peptides, and analogues, Amino acids and derivatives, Alpha amino acids and derivatives, Thiodioxopiperazines, Indoles and derivatives, N-methylpiperazines, Tertiary carboxylic acid amides, Pyrrolidines, Secondary alcohols, Lactams, Azacyclic compounds, Primary alcohols, Organonitrogen compounds, Organic oxides, Hydrocarbon derivatives, Carbonyl compounds |
| Hexadehydroastechrome | 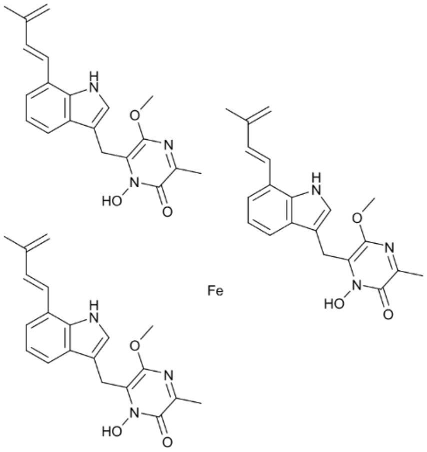 | **Organoheterocyclic compounds**, Indoles and derivatives, Indoles, 3-alkylindoles, Styrenes, Methoxypyrazines, Alkyl aryl ethers, Substituted pyrroles, Heteroaromatic compounds, Lactams, Organic transition metal salts, Azacyclic compounds, Organopnictogen compounds, Organonitrogen compounds, Organic oxides, Hydrocarbon derivatives |
| Isochaetominine | 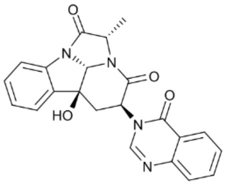 | **Organoheterocyclic compounds**, Indoles and derivatives, Pyridoindoles, Pyridoindolones, Alpha carbolines, Quinazolines, Alpha amino acids and derivatives, Indoles, Imidazopyridines, Pyrimidones, Piperidinones, Delta lactams, Pyridines and derivatives, Imidazolidinones, Benzenoids, Tertiary carboxylic acid amides, Tertiary alcohols, Heteroaromatic compounds, Azacyclic compounds, Organopnictogen compounds, Organonitrogen compounds, Organic oxides, Hydrocarbon derivatives, Carbonyl compounds |
| Neosartorin | 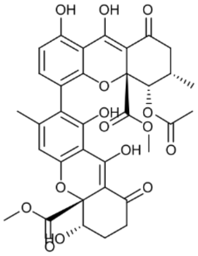 | **Organoheterocyclic compounds**, Benzopyrans, 1-benzopyrans, Dibenzopyrans, Xanthenes, Tricarboxylic acids and derivatives, Beta hydroxy acids and derivatives, Alkyl aryl ethers, 1-hydroxy-4-unsubstituted benzenoids, 1-hydroxy-2-unsubstituted benzenoids, Vinylogous acids, Methyl esters, Secondary alcohols, Ketones, Cyclic alcohols and derivatives, Polyols, Oxacyclic compounds, Enols, Organic oxides, Hydrocarbon derivatives |
| Patientoside A | 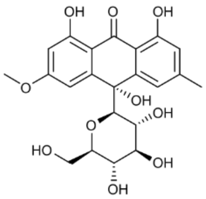 | **Benzenoids,** Anthracenes, Hexoses, C-glycosyl compounds, Anisoles, Aryl ketones, 1-hydroxy-2-unsubstituted benzenoids, 1-hydroxy-4-unsubstituted benzenoids, Alkyl aryl ethers, Oxanes, Vinylogous acids, Tertiary alcohols, Secondary alcohols, Polyols, Oxacyclic compounds, Dialkyl ethers, Organic oxides, Hydrocarbon derivatives, Primary alcohols |
| Pyripyropene B | 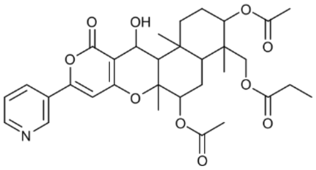 | **Lipids and lipid-like molecules**, Steroids and steroid derivatives, Hydroxysteroids, 1-hydroxysteroids, Naphthopyrans, Naphthalenes, Tricarboxylic acids and derivatives, Alkyl aryl ethers, Pyranones and derivatives, Pyridines and derivatives, Vinylogous esters, Heteroaromatic compounds, Secondary alcohols, Lactones, Carboxylic acid esters, Oxacyclic compounds, Azacyclic compounds, Organic oxides, Hydrocarbon derivatives, Carbonyl compounds, Organonitrogen compounds, Organopnictogen compounds |

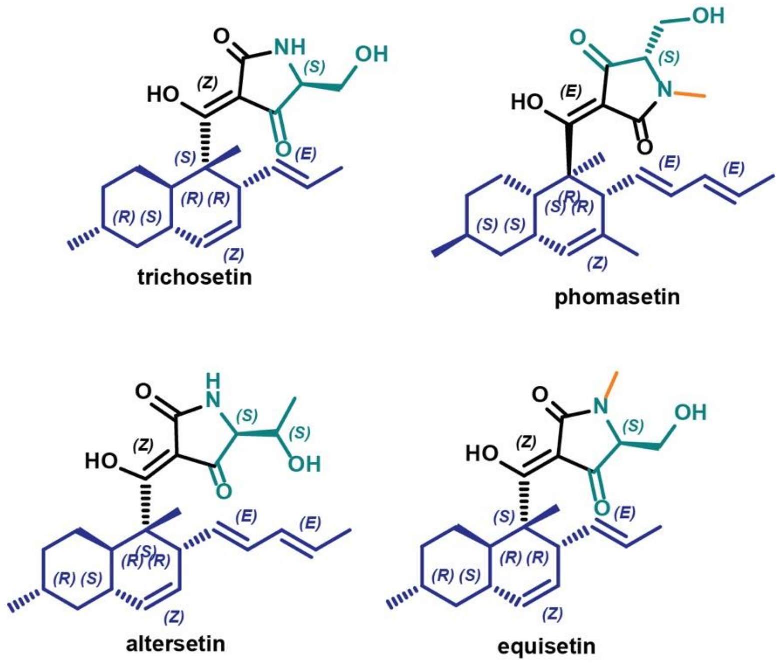


**Fig. S11.** Compounds from the equisetin structural class that have associated known gene clusters. The scaffold includes a hydrocarbon decalin core varying in methyl and alkenyl substituents and stereochemistry. A tetramic acid moiety derived from serine or threonine is conjugated to the decalin core. N-methylation of the tetramic acid amide is present in equisetin and phomasetin.

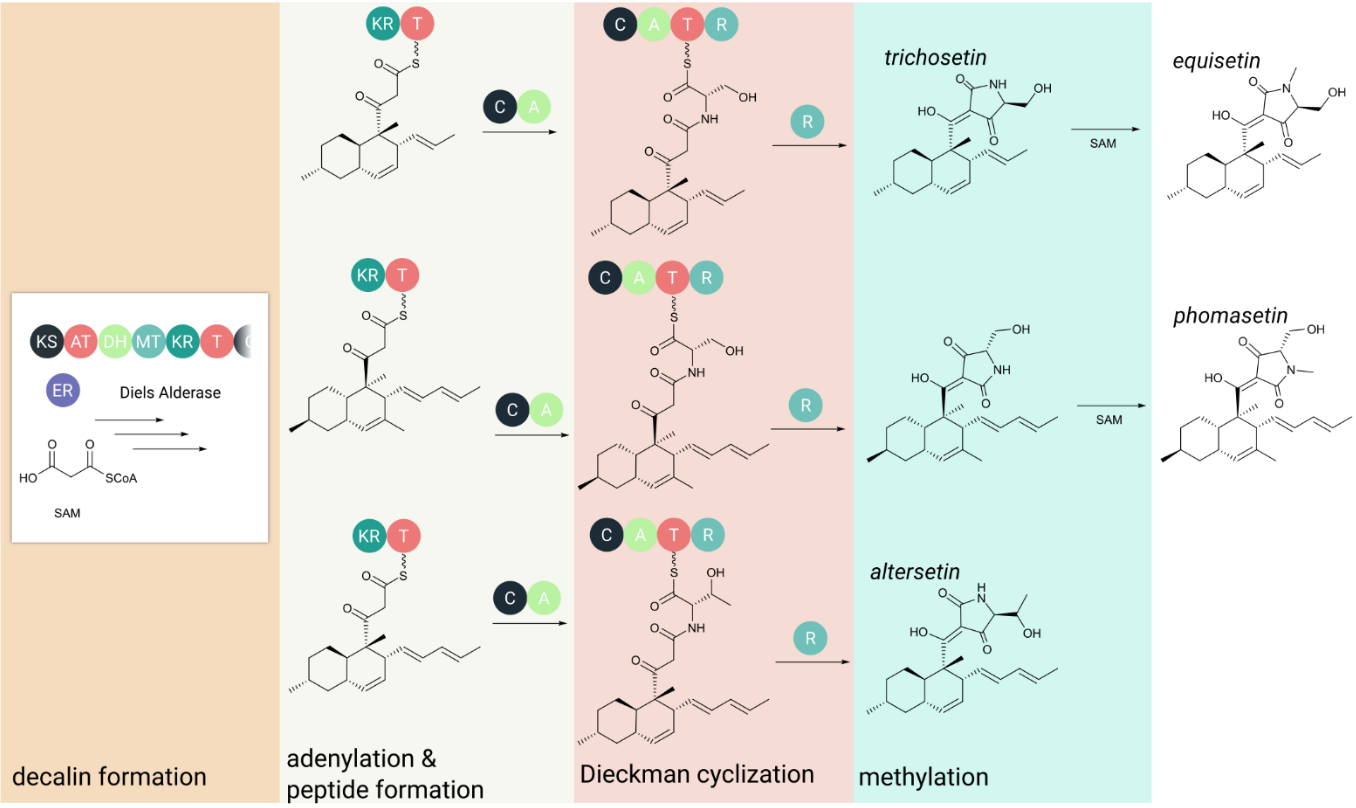


**Fig. S12.** The biosynthetic pathway for equisetin and related compounds. First the core decalin ring is constructed by a hybrid nonribosomal peptide synthetase-polyketide synthase (NRPS-PKS) enzyme. The PKS domains within the backbone enzyme act in an iterative fashion typical of fungal PKS enzymes, assembling the decalin core from malonyl-CoA monomers. This step is supplemented by the action of a standalone enoyl reductase for ketide monomer reduction and a Diels-Alderase that directs ring closure and controls stereochemistry (14, 15). Second, an NRPS module condenses an amino acid to the decalin core (16). A terminal reductase domain catalyzes Dieckman cyclization to release the intermediate as a tetramic acid, the third step (17). In the final pathway step, a methyltransferase catalyzes N-methylation of the tetramic acid amide (16).

**Table S3**. The 90 gene cluster families (from total n = 12,067) that are exceptional in that they span multiple taxonomic classes. The Reference column indicates a single GenBank accession number and organism for the backbone enzyme. In cases of multiple backbone enzymes, the provided GenBank reference corresponds to the backbone enzyme in bold text. Abbreviations are as follows: DHONTB, dihydroxy-6-[(3E,5E,7E)-2-oxonona-3,5,7-trienyl]-benzaldehyde; HAS, hexadehydroastechrome; KS, ketosynthase, AT, acyltransferase; DH, dehydratase; ER, enoyl reductase; KR, ketoreductase; MT, methyltransferase; SAT, starter acyltransferase; PT, product template; A, adenylation; T, thiolation; R, reductase; C, condensation; ICS, isocyanide synthase; DMAT, dimethylallyltransferases; NRPS, nonribosomal peptide synthetase; PKS, polyketide synthase; HRPKS, highly reducing polyketide synthase; NRPKS, nonreducing polyketide synthase; E, Eurotiomycetes; L, Leotiomycetes; S, Sodariomycetes; D, Dothidiomycetes; X, Xylonomycetes; LEC, Lecanoromycetes.

| GCF | REFERENCE | BACKBONE | TAXONOMIC CLASSES |
|---|---|---|---|
| **HRPKS_30 (DHONTB)** | *Aspergillus nidulans FGSC A4* (CBF86052) | **PKS (KS-AT-DH-ER-KR-T)**, PKS (SAT-KS-AT-PT-T-R) | E, S |
| **NRPKS_1343 (BIKAVERIN)** | *Fusarium fujikuroi* (SCO46930.) | PKS (KS-AT-PT-T-TE) | L, S |
| **NRPKS_791 (MELANIN)** | *Cadophora sp. DSE1049* (PVH73815) | PKS (SAT-KS-AT-PT-T-T-TE) | D, L, S |
| **NRPS_607 (CHAETOGLOBOSIN)** | *Aspergillus lentulus* (GAQ05296) | PKS (KS-AT-DH-MT-ER-KR-T-C-A-T-R) | D, E |
| **NRPS_63 (CHRYSOGINE)** | *Aspergillus arachidicola* (PIG85941) | NRPS (C-A-T-C-A-T-C) | E, S |
| **NRPKS_375 (CONIDIAL YELLOW PIGMENT)** | *Aspergillus sydowii CBS 593.65* (OJJ57401) | PKS (SAT-KS-AT-PT-T) | E, L, LEC, S |
| **NRPS_690 (CYTOCHALASIN)** | *Aspergillus clavatus NRRL 1* (EAW09117) | PKS (KS-AT-DH-MT-ER-KR-T-C-A-T-R) | E, L, S |
| **NRPS_138 (EQUISETIN)** | *Alternaria alternata* (OWY46706) | PKS (KS-AT-DH-MT-ER-KR-T-C-A-T-R) | D, E, S |
| **NRPS_123 (FUMITREMORGIN)** | *Aspergillus fumigatus* (OXN23238) | NRPS (A-T-C-A-T-C) | E, L, S |
| **NRPS_1705 (FUMONISIN)** | *Fusarium verticillioides* (RBR13858) | PKS (KS-AT-DH-MT-ER-KR-T) | D, S |
| **NRPS_442 (HAS)** | *Aspergillus fumigatus* (OXN25028) | NRPS (A-T-C-A-T-C-T) | E, S |
| **NRPKS_147 (MELANIN)** | *Alternaria alternata* (OAG24502) | PKS (SAT-KS-AT-PT-T-T-TE) | D, E |
| **NRPS_101 (PHOMASETIN)** | *Aspergillus clavatus NRRL 1* (EAW07624) | PKS (KS-AT-DH-MT-ER-KR-T-C-A-T-R) | D, E, S, X |
| **NRPS_1149 (SERINOCYCLIN)** | *Metarhizium acridum CQMa 102* (EFY85053) | NRPS (A-T-C-C-A-T-C-A-T-C-A-T-C-C-A-T-C-C-A-T-C-C-A-T-C) | L, S |
| **HYBRIDS_151 (SWAINSONINE)** | *Clohesyomyces aquaticus* (ORY11783) | PKS (A-T-KS-AT-KR-T-R) | E, S |
| **NRPS_2042 (UCS1025A)** | *Oidiodendron maius Zn* (KIM94019) | PKS (KS-AT-DH-MT-ER-KR-T-C-A-T-R) | L, S |
| **DMAT_140** | *Ophiocordyceps australis* (PHH64516) | DMAT | E, S |
| **DMAT_401** | *Colletotrichum orchidophilum* (OHF04557) | PKS (SAT-KS-AT-MT-PT-T-TE), DMAT | L, S |
| **DMAT_411** | *Cadophora sp. DSE1049* (PVH84683) | DMAT | L, S |
| **HRPKS_1152** | *Meliniomyces bicolor E* (PMD61012) | PKS (KS-AT-DH-MT-ER-KR-T) | L, S |
| **HRPKS_128** | *Pezoloma ericae* (PMD17755) | PKS (KS-AT-DH-MT-ER-KR-T) | E, L |
| **HRPKS_1289** | *Acremonium chrysogenum ATCC 11550* (KFH46614) | PKS (KS-AT-DH-MT-ER-KR-T-Carnitine_acyltransferase) | L, S |
| **HRPKS_1318** | *Colletotrichum higginsianum IMI 349063* (OBR06526) | **PKS (KS-AT-DH-MT-ER-KR-T-C)**, PKS (KS-AT-DH-MT-ER-KR-T) | L, S |
| **HRPKS_159** | *Penicillium griseofulvum* (KXG49005) | **PKS (SAT-KS-AT-PT-MT-R)**, PKS (KS-AT-DH-MT-ER-KR-T) | E, S |
| **HRPKS_170** | *Penicillium camemberti* (CRL31088) | PKS (KS-AT-DH-MT-ER-KR-T) | E, L |
| **HRPKS_216** | *Aspergillus sydowii CBS 593.65* (OJJ61536) | NRPS (C-A-T-C-C-A-T-C-A-T-C-A-T-C) | E, L |

| | | | |
|---|---|---|---|
| **HRPKS_495** | *Aspergillus uvarum CBS 121591* (PYH83208) | PKS (KS-AT-DH-ER-KR-T) | E, S |
| **HRPKS_53** | *Colletotrichum chlorophyti* (OLN93260) | PKS (KS-AT-DH-ER-KR-T-R) | E, S |
| **HRPKS_597** | *Cordyceps sp. RAO-2017* (PHH90746) | PKS (KS-AT-DH-ER-KR-T) | E, S |
| **HRPKS_678** | *Pseudogymnoascus sp. VKM F-3557* (KFX86927) | PKS (KS-AT-DH-MT-ER-KR-T-Carnitine_acyltransferase) | E, L |
| **HRPKS_694** | *Phialocephala subalpina* (CZR67900) | PKS (KS-AT-DH-MT-ER-KR-T) | E, L |
| **HRPKS_882** | *Fusarium fujikuroi* (SCN83763) | PKS (KS-AT-DH-MT-ER-KR-T) | S, X |
| **HYBRIDS_195** | *Aspergillus ochraceoroseus IBT 24754* (PTU20620) | PKS (KS-AT-DH-MT-ER-KR-T-C-A-T-R) | E, S |
| **HYBRIDS_215** | *Penicillium camemberti* (CRL19370) | PKS (KS-AT-DH-MT-ER-KR-T) | E, S |
| **HYBRIDS_506** | *Talaromyces stipitatus ATCC 10500* (EED18841) | PKS (KS-AT-DH-MT-ER-KR-T) | E, L |
| **HYBRIDS_9** | *Penicillium subrubescens* (OKP00032) | PKS (KS-AT-DH-MT-ER-KR-T-C-A-T-R) | D, E, L |
| **NRPKS_1290** | *Pseudogymnoascus sp. 05NY08* (OBT71831) | PKS (KS-AT-DH-MT-ER-KR-T-C) | E, L |
| **NRPKS_1320** | *Coniochaeta pulveracea* (RKU46359) | PKS (KS-AT-DH-ER-KR-T) | E, S |
| **NRPKS_1782** | *Phialocephala scopiformis* (KUJ09200) | **PKS (SAT-KS-AT-PT-T-T-TE)**, PKS (DH-KR) | D, L |
| **NRPKS_1988** | *Pseudogymnoascus sp. 23342-1-I1* (OBT65120) | PKS (SAT-KS-AT-PT-T) | D, L |
| **NRPKS_20** | *Pseudogymnoascus sp. VKM F-103* (KFY80205) | PKS (KS-AT-DH-MT-ER-KR-T-C-A-T-R) | |
| **NRPKS_250** | *Aspergillus lentulus* (GAQ09994) | PKS (KS-AT-DH-ER-KR-T) | E, L |
| **NRPKS_437** | *Aspergillus kawachii IFO 4308* (GAA83965) | PKS (A-T-KS-AT-KR-T-R) | |
| **NRPKS_447** | *Endocarpon pusillum Z07020* (ERF68696) | PKS (A-T-KS-AT-KR-T-R) | |
| **NRPKS_5** | *Penicillium nalgiovense* (OQE96240) | PKS (SAT-KS-AT-PT-T) | E, X |
| **NRPKS_510** | *Trichoderma asperellum CBS 433.97* (PTB35070) | PKS (SAT-KS-AT-PT-T) | E, S |
| **NRPKS_548** | *Fusarium oxysporum f. sp. cepae* (RKK07595) | PKS (KS-AT-DH-MT-KR-T-C-A-T-R) | |
| **NRPKS_604** | *Scedosporium apiospermum* (KEZ41293) | PKS (KS-AT-DH-MT-KR-T-R) | E, S |
| **NRPKS_787** | *Penicillium griseofulvum* (KXG49279) | PKS (KS-AT-DH-MT-ER-KR-T-R) | D, E, L |
| **NRPS_1018** | *Pseudogymnoascus sp. VKM F-3808* (KFX99775) | NRPS (A-T-C-A-T-R) | D, L |
| **NRPS_1055** | *Bipolaris victoriae FI3* (EUN25091) | PKS (KS-AT-DH-MT-ER-KR-T-C-A-T-R) | D, L |
| **NRPS_1064** | *Coleophoma cylindrospora* (RDW81833) | **PKS (SAT-KS-AT-PT-T-T-TE)**, NRPS (C-A-T) | E, L |
| **NRPS_111** | *Aspergillus brasiliensis CBS 101740* (OJJ75537) | **PKS (KS-AT-DH-MT-ER-KR-T-C-A-T-R)**, NRPS (A-T-C) | E, S |
| **NRPS_1222** | *Talaromyces stipitatus ATCC 10500* (EED13058) | **PKS (KS-AT-DH-MT-ER-KR-T)**, NRPS (A-T-C-A-T-C-T) | E, S |
| **NRPS_1295** | *Penicillium steckii* (OQE21884) | PKS (KS-AT-DH-MT-ER-KR-T-C-A-T-R) | E, L, S |
| **NRPS_1301** | *Aspergillus bombycis* (OGM48141) | PKS (KS-AT-DH-ER-KR-T-C-A-T-R) | E, S |
| **NRPS_1372** | *Fusarium avenaceum* (KIL86455) | PKS (KS-AT-DH-MT-ER-KR-T-C-A-T-R) | E, S |
| **NRPS_1410** | *Helicocarpus griseus UAMH5409* (PGH19023) | PKS (KS-AT-DH-MT-ER-KR-T-C-A-T-R) | E, S |
| **NRPS_1417** | *Aspergillus heteromorphus CBS 117.55* (PWY81896) | PKS (KS-AT-DH-MT-ER-KR-T-C-A-T-R) | E, L |
| **NRPS_151** | *Madurella mycetomatis* (KXX75968) | PKS (KS-AT-DH-MT-ER-KR-T-C-A-T-R) | |
| **NRPS_1545** | *Fusarium avenaceum* (KIL87829) | **PKS (KS-AT-DH-ER-KR-T)**, NRPS (T-C-A-T-C-A-T-C-A-T-C-T) | E, L, S |
| **NRPS_1559** | *Pseudogymnoascus sp. VKM F-3775* (KFY27678) | PKS (KS-AT-DH-MT-ER-KR-T-C-A-T-R) | E, L |
| **NRPS_1586** | *Metarhizium rileyi RCEF 4871* (OAA34246) | NRPS (A-T-C-A-T-C-A-T-R) | E, S |
| **NRPS_2023** | *Colletotrichum graminicola M1.001* (EFQ35223) | PKS (KS-AT-DH-MT-ER-KR-T-C-A-T-R) | D, S |

| | | | |
|---|---|---|---|
| **NRPS_2636** | *Bipolaris sorokiniana ND90Pr* (EMD59100) | NRPS (A-T-C) | E, S |
| **NRPS_283** | *Aspergillus bombycis* (OGM44044) | PKS (KS-AT-DH-ER-KR-T-C-A-T-R) | E, S |
| **NRPS_353** | *Beauveria bassiana ARSEF 2860* (EJP61198) | PKS (KS-AT-DH-MT-KR-T-C-A-T-R) | E, S |
| **NRPS_41** | *Aspergillus steynii IBT 23096* (PLB43453) | NRPS (A-T-C-A-T-C) | E, L |
| **NRPS_457** | *Coleophoma crateriformis* (RDW59260) | **PKS (KS-AT-DH-MT-ER-KR-T)**, NRPS (A-T-C) | E, L |
| **NRPS_480** | *Capronia coronata CBS 617.96* (EXJ78804) | NRPS (A-T-C-T-C) | E, L |
| **NRPS_514** | *Aspergillus mulundensis* (RDW86494) | **PKS (KS-AT-DH-MT-ER-KR-T)**, NRPS (T-C-A-T-C-A-T-C-A-T-C-A-T-C) | E, S, L |
| **NRPS_569** | *Cladophialophora carrionii* (OCT48933) | NRPS (A-T-C-A-T-C-T-C-A-T-C-T-C-T-C) | E, L |
| **NRPS_648** | *Hypoxylon sp. CO27-5* (OTA94984) | **PKS (KS-AT-DH-MT-ER-KR-T-C-A-T-R)**, NRPS (A-T-R) | E, S |
| **NRPS_777** | *Cordyceps fumosorosea ARSEF 2679* (OAA69787) | PKS (KS-AT-DH-MT-ER-KR-T-C-A-T-R) | E, S |
| **NRPS_871** | *Aspergillus fischeri NRRL 181* (EAW20390) | **NRPS (A-T-C-A-T-R)**, NRPS (A-T-C-A-T-C) | D, E, L, S |
| **NRPS_932** | *Aspergillus costaricaensis CBS 115574* (RAK83302) | **PKS (KS-AT-DH-MT-ER-KR-T)**, PKS (KS-AT-DH-MT-ER-KR-T) | D, E, L |
| **NRPSLIKE_10** | *Aspergillus ochraceoroseus* (KKK21469) | NRPS-like (ICS-A-T-Transferase) | D, E, S |
| **NRPSLIKE_1029** | *Cladophialophora carrionii CBS 160.54* (ETI26263) | NRPS-like (A-T-R) | E, L |
| **NRPSLIKE_11** | *Aspergillus lentulus* (GAQ04120) | NRPS-like (ICS-A-T-Transferase) | E, S |
| **NRPSLIKE_1277** | *Amorphotheca resinae ATCC 22711* (PSS07172) | NRPS-like (A-T-R) | L, S |
| **NRPSLIKE_128** | *Exophiala oligosperma* (KIW43198) | NRPS-like (A-T-Transferase) | D, S |
| **NRPSLIKE_1465** | *Neonectria ditissima* (KPM46454) | NRPS-like (A-T-R) | D, S |
| **NRPSLIKE_1739** | *Ophiocordyceps australis* (PHH75740) | NRPS-like (A-T-TE) | D, L, S |
| **NRPSLIKE_22** | *Cladophialophora bantiana CBS 173.52* (KIW93789) | NRPS-like (A-T-R-DH) | |
| **NRPSLIKE_266** | *Penicillium occitanis* (PCG97091) | **NRPS-like (A-T-R)**, NRPS-like (A-T-R) | E, L |
| **NRPSLIKE_869** | *Cladophialophora bantiana CBS 173.52* (KIW89508) | NRPS-like (A-T-R) | E, L |
| **NRPSLIKE_873** | *Cladophialophora carrionii CBS 160.54* (ETI24620) | NRPS-like (A-T-R) | E, L |
| **NRPSLIKE_899** | *Talaromyces marneffei ATCC 18224* (EEA18553) | NRPS-like (A-T-R) | E, L |
| **TERPENE_1140** | *Exserohilum turcica Et28A* (EOA88708) | trichodiene synthase | D, L |
| **TERPENE_139** | *Penicillium camemberti* (CRL18805) | terpene cyclase | E, S |

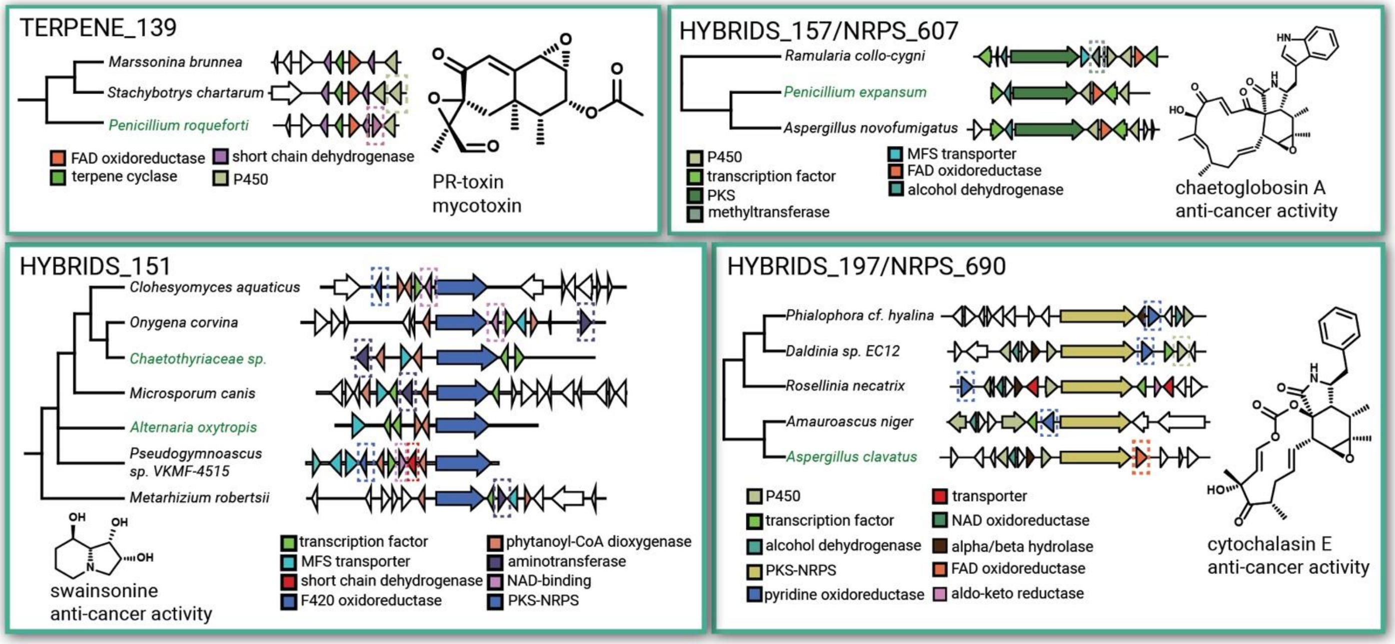


**Fig. S13.** Diversification of chemical scaffolds across gene cluster families. The GCF for PR-toxin (TERPENE_139), a DNA polymerase mycotoxin produced by *Penicillium roqueforti* (18), contains an additional P450 enzyme in a BGC from the Sordariomycete *Stachybotrys chartarum*. The GCF for chaetoglobosin A, a scaffold with a variety of anti-cancer activities (19), contains a methyltransferase in a BGC from the Dothideomycete *Ramularia collo-cygni* not present in the experimentally-characterized BGC from *Penicillium expansum*. The GCF for swainsonine (HYBRIDS_151), an α-mannosidase inhibitor advanced to clinical trials as a potential anti-cancer therapeutic (20, 21), contains variable F420 oxidoreductase, short chain dehydrogenase, and an NAD oxidoreductase, and aminotransferase enzymes. In the GCF for cytochalasin E (HYBRIDS_197), a compound with anticancer activity, BGCs differ in the presence/absence of a pyridine oxidoreductase and an FAD oxidoreductase present in the experimentally-characterized *Aspergillus clavatus* BGC.

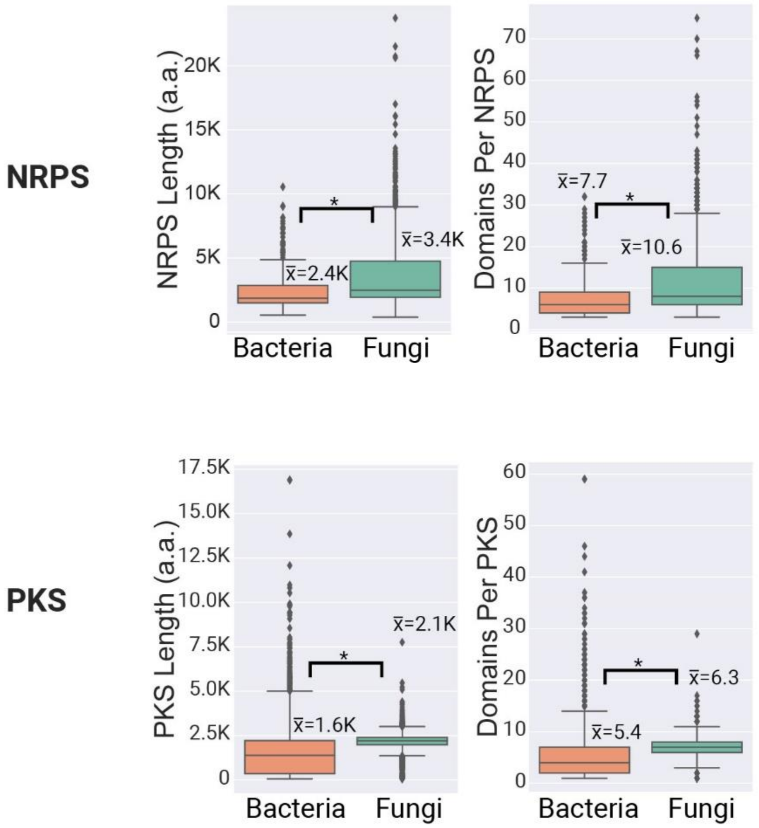


**Fig. S14.** Comparison of fungal and bacterial NRPS and PKS backbone sizes. For both NRPS and PKS enzymes, fungal backbones are longer both in terms of amino acids and catalytic domains per backbone enzyme.

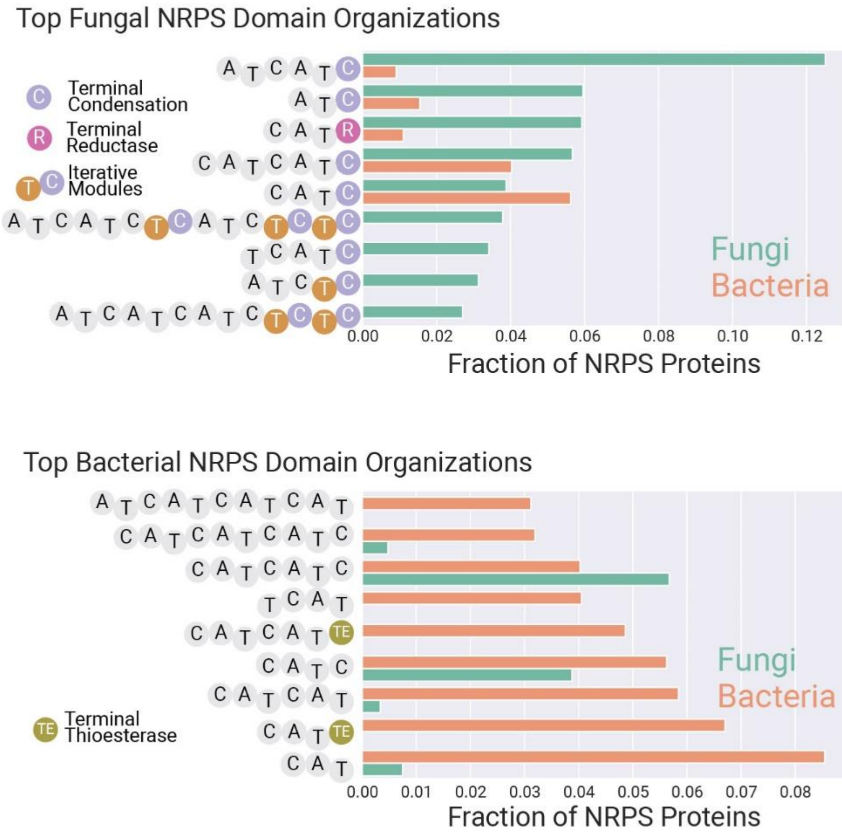


**Fig. S15.** Comparison of fungal and bacterial NRPS domain organizations. In fungi (top), the most common NRPS domain organizations include terminal condensation or thioester reductase domains. Fungal NRPS enzymes also commonly employ iterative modules. In bacteria, the most common NRPS domain organizations feature terminal thioesterase domains and/or N-terminal condensation domains that interact with an upstream NRPS enzyme catalyze N-acylation.

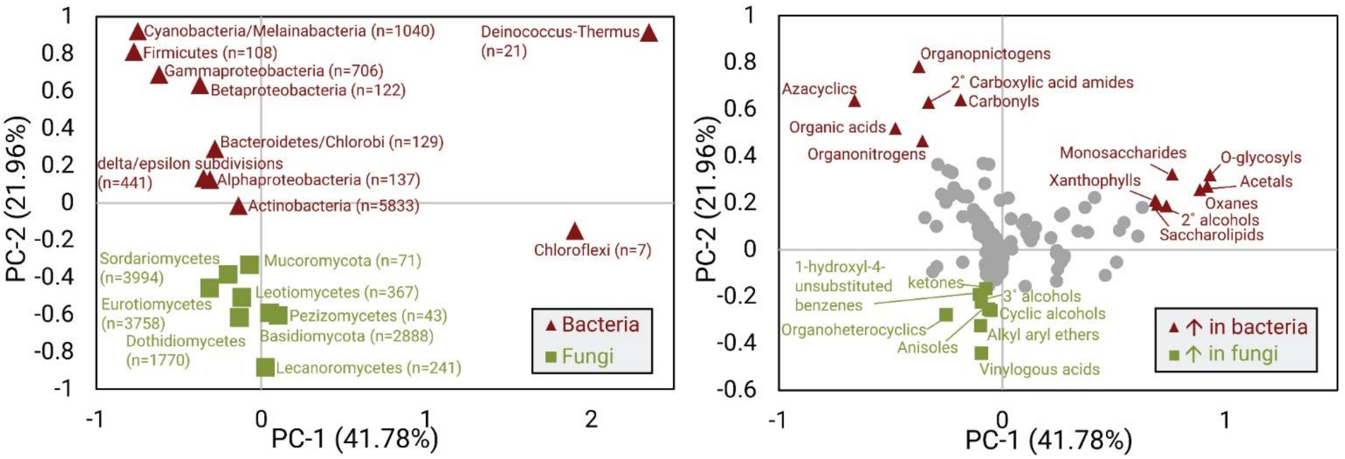


**Fig. S16**. PCA plot (left) and associated loadings plot (right) of bacterial and fungal chemical space. Fungal and bacterial taxonomic groups represent distinct regions in this space. Fungi are distinguished from bacteria due to an increased frequency of chemical ontology terms associated with aromatic polyketides, such as anisoles, ketones, and alkyl aryl ethers. Bacteria are distinguished largely due to peptide-associated chemical ontology terms (i.e. organic acids, azacyclics, amides).

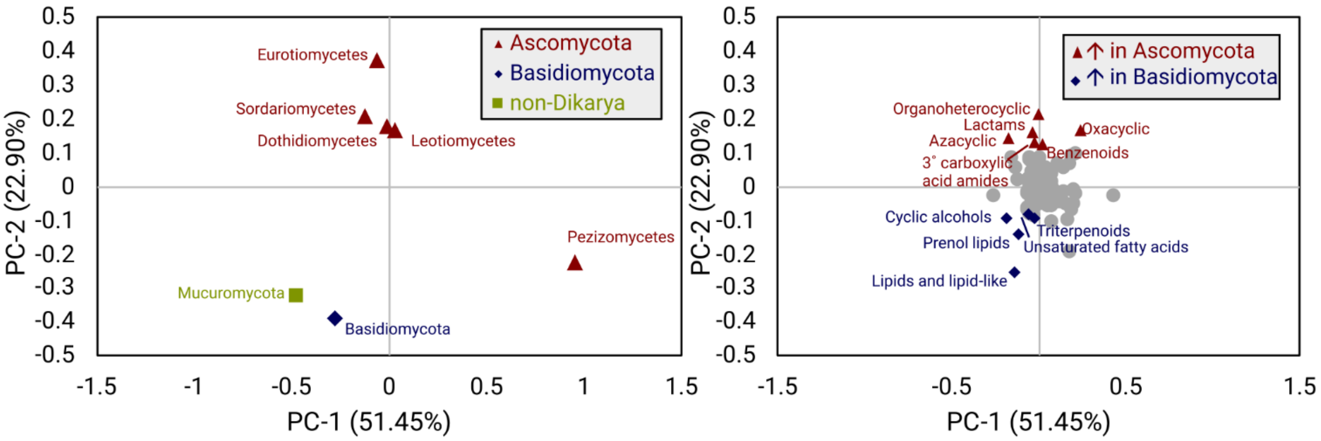


**Fig. S17**. PCA analysis of fungal chemical space. Eurotiomycetes, Sordariomycetes, Dothideomycetes, and Leotiomycetes (Ascomycota) are distinct largely based on polyketide and peptide-related chemical ontology terms, such as *azacyclic*, *Oxacyclic*, *Benzenoids*, and *Lactams*. Lipid-associated chemical ontology terms are prevalent in Basidiomycota and Mucoromycota.

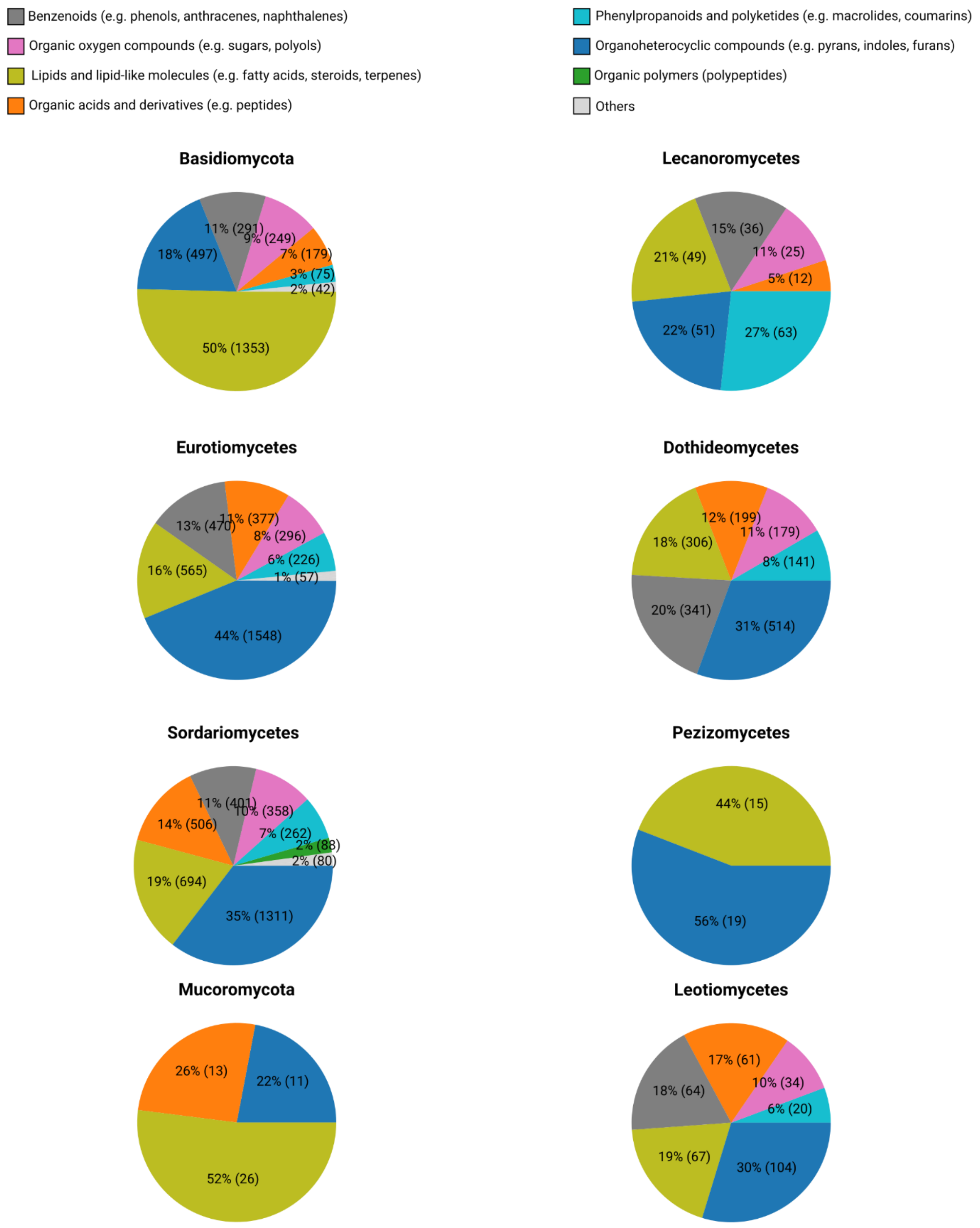


**Fig. S18**. Breakdown of chemical superclasses in fungal taxa. The chemical space of distinct fungal taxonomic groups varies dramatically. Basidiomycota and Mucoromycota are both ~50% lipids. Other taxonomic groups contain a higher fraction of organoheterocyclic compounds.

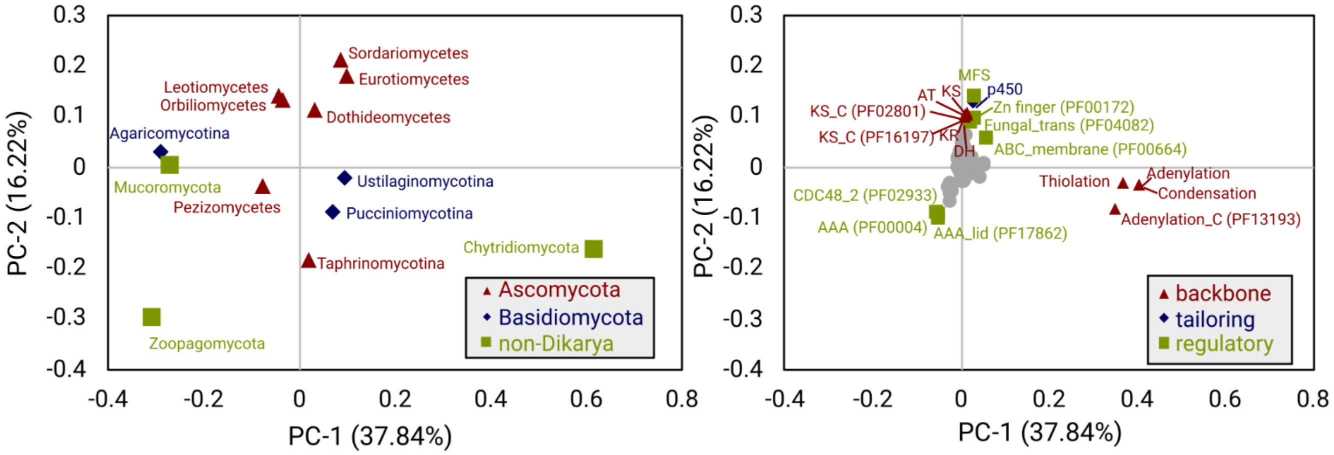


**Fig. S19**. PCA plot (left) and associated loading plot (right) of biosynthetic domains contained within NRPS-containing biosynthetic gene clusters. Chytridiomycota are pulled in the positive direction on the x-axis due to their high frequency of large NRPS backbone enzymes containing many adenylation, condensation, and thiolation domains, while Pezizomycotina are largely pulled in the "up" direction due to the presence of NRPS-PKS hybrids.

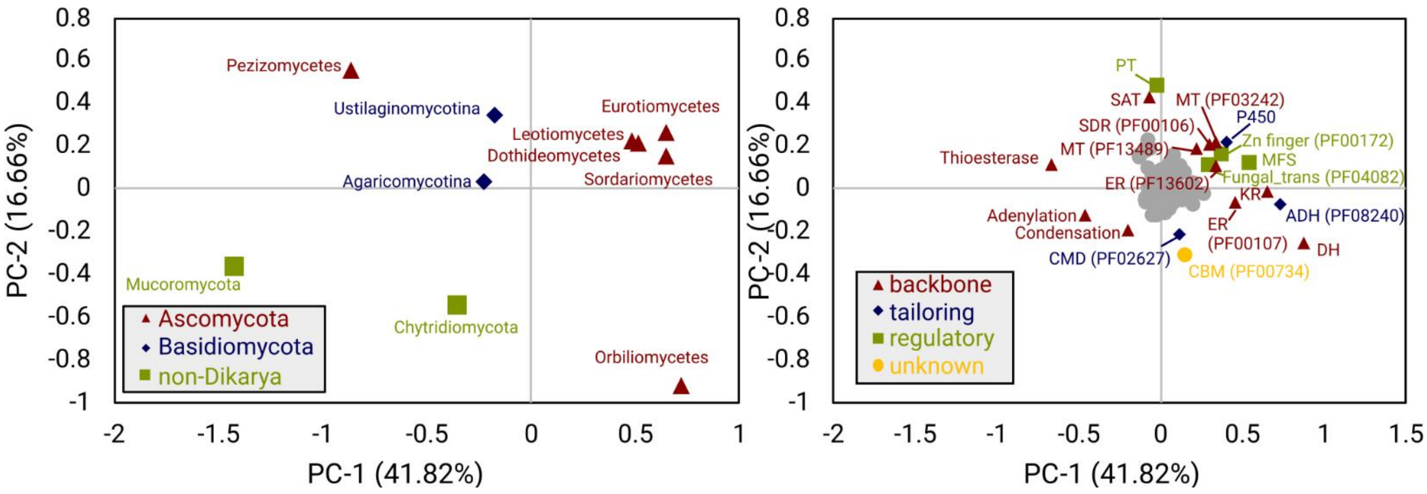


**Fig. S20**. PCA plot (left) and associated loading plot (right) of biosynthetic domains contained within PKS-containing biosynthetic gene clusters. Eurotiomycetes, Leotiomycetes, Dothideomycetes, and Sordariomycetes contain the most PKS backbone enzymes, and are pulled to the right by the corresponding PKS domains. Several regulatory elements are associated with these backbone genes, providing insight into the way fungi regulate PKS biosynthesis.

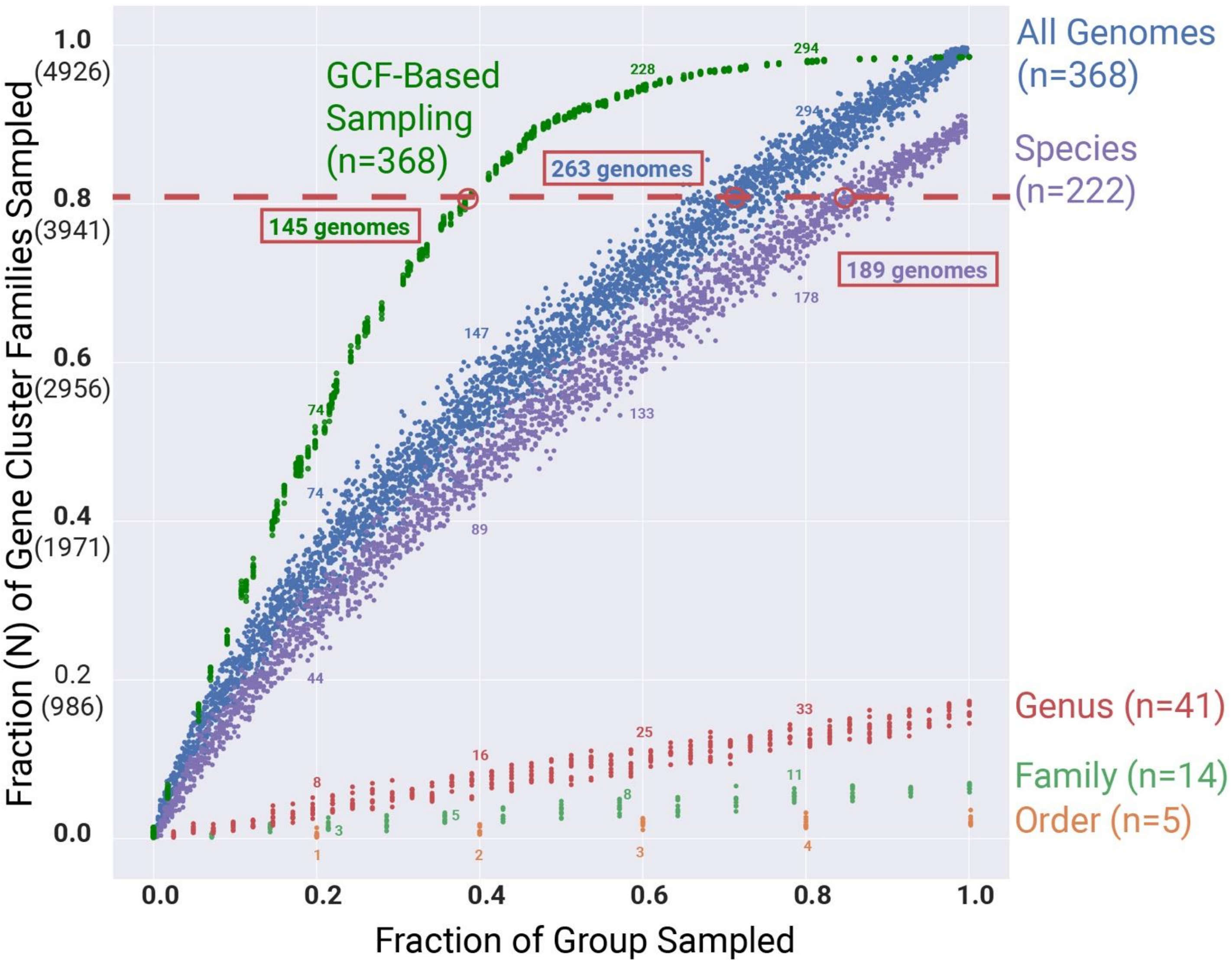


**Fig. S21**. A roadmap for sampling Eurotiomycetes genomes for natural products discovery based on shared GCFs. Each curve shows the fraction of Eurotiomycetes GCFs that would be present in genomes sampled using different approaches. *All Genomes* shows the results of randomly sampling from all 368 Eurotiomycetes genomes. *Species* and other taxonomic ranks shows the result of randomly sampling unique species, genera, families, or orders. *GCF-Based Sampling* shows the result of sampling clusters of organisms that share GCFs ("clusters" representing the results of density-based clustering, not *biosynthetic gene clusters*). The red boxed numbers indicate the number of genomes required reach 80% GCF coverage, the threshold indicated by the dashed red line. Small numbers along each curve indicate the number of genomes randomly sampled from each group. GCF-based sampling of organisms reaches 80% coverage of GCFs after 145 genomes sampled, species-based sampling of organisms requires 189 genomes, and random sampling of all genomes requires 263 genomes to reach this threshold. This indicates that sampling of organisms for biosynthetic pathway and compound discovery based on GCF overlap can provide a more efficient means of accessing these GCFs. Each random sampling of genomes was performed using 1000 iterations.

.

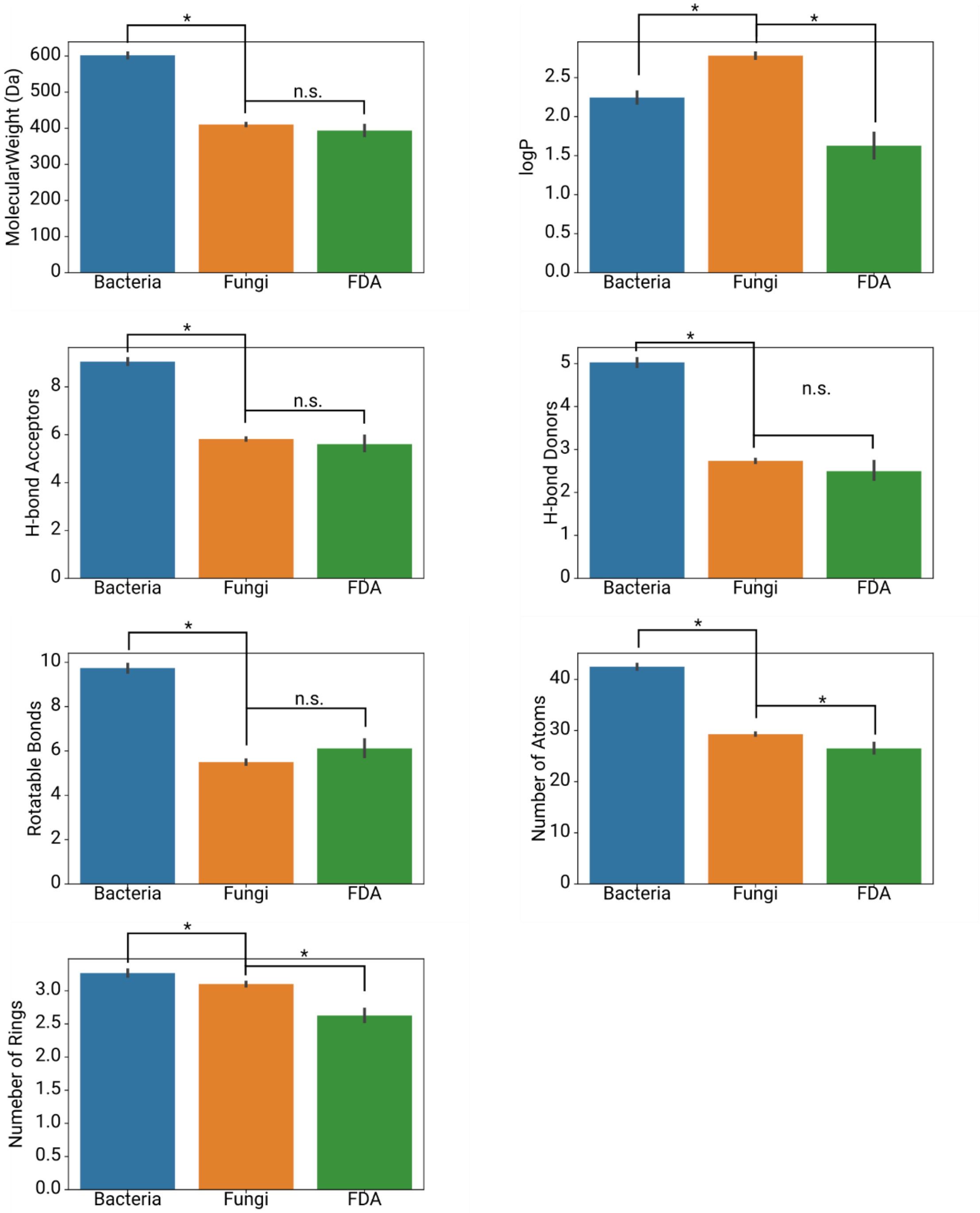


**Fig. S22**. Comparison of the pharmacological properties of bacterial (n=9,382), fungal (n=15,213), and FDA-approved compounds (n=2884). Error bars represent 95% confidence intervals determined by bootstrap sampling. Asterisks indicate statistically-significant differences between the means ($p < 0.01$, Student's t-test).

| BGC TYPE | DOMAINS PRESENT | DOMAINS ABSENT |
|---|---|---|
| **NRPS** | Adenylation, condensation | N/A |
| **HR-PKS** | Ketosynthase, dehydratase | N/A |
| **NR-PKS** | Ketosynthase and product template or starter acyltransferase | N/A |
| **HYBRID NRPS-PKS** | Adenylation, ketosynthase | N/A |
| **NRPS-LIKE** | Adenylation | Condensation |
| **DMAT** | Dimethylallyl transferase | N/A |
| **TERPENE** | Terpene synthase, terpene cyclase, trichodiene synthase, or polyprenyl synthetase | N/A |

**Table S4**. Protein domain rules for classifying gene clusters as nonribosomal peptide synthase (NRPS), highly-reducing polyketide synthase (HR-PKS), nonreducing polyketide synthase (NR-PKS) hybrid NRPS-PKS, NRPS-like, dimethylallyl transferase (DMAT), or terpene.
